# Lateral organization of cytochrome *b_6_f* in thylakoid membranes controls photosynthetic electron transfer efficiency

**DOI:** 10.64898/2026.08.18.745480

**Authors:** Afifa Zaeem, Davide Tamborrini, Martin Scholz, Wojciech Wietrzynski, Markus Schwarzländer, Benjamin D. Engel, Michael Hippler, Felix Buchert

## Abstract

Photosynthetic electron transfer relies on the coordinated function and spatial organization of large protein complexes within the thylakoid membrane. The cytochrome *b_6_f* complex (*b_6_f*) functionally interconnects photosystem (PS) II and PSI in photosynthetic electron transfer and is equally distributed between appressed and non-appressed thylakoid membranes. Here, we investigate the functional link between the lateral distribution of *b_6_f* and efficient photosynthetic electron flow in *Chlamydomonas reinhardtii*. We engineered strains with stromal fusions between PetA of *b_6_f* and two fluorescent proteins (FPs) of different molecular mass: Clover and ATeam. Under oxic conditions, these strains exhibited significantly slower electron transfer rates (ETR), lower PSII quantum yields, and increased donor-side limitation of PSI. State transitions were diminished in the fusion strains, accompanied by a strong impairment of STT7-dependent function, suggesting that the presence of fused FPs at *b_6_f* interfere with STT7 function. Yet, ETR phenotypes were STT7-independent and FP fusion did not impact intrinsic *b_6_f* function. In situ cryogenic electron tomography revealed a significant depletion of *b_6_f* from appressed thylakoid membranes in the ATeam strains, while the overall membrane protein concentration remained unchanged. Overall, our data indicate that a balanced distribution of *b_6_f* between appressed and non-appressed thylakoid membranes is essential for regulating photosynthetic electron transfer, highlighting the functional importance of thylakoid molecular architecture in vivo.

## Introduction

Photosynthesis converts light energy into chemical energy through the coordinated action of photosystem II (PSII) and photosystem I (PSI), each associated with its respective light-harvesting complexes (LHCs), LHCII and LHCI. Light-driven charge separation at PSII initiates electron transfer through the thylakoid membrane to PSI, where electrons reduce ferredoxin. In linear electron flow (LEF), ferredoxin donates electrons to ferredoxin–NADP(H) oxidoreductase, producing NADPH for CO_2_ fixation. Simultaneously, proton-coupled electron transport drives ATP synthesis by establishing a proton motive force composed of chemical proton gradient (ΔpH) and electric field (ΔT) across the thylakoid membrane.

The cytochrome *b_6_f* complex (*b_6_f*), a major regulatory hub, mediates electron transfer between PSII and PSI by oxidizing plastoquinol (PQH_2_) and reducing plastocyanin (PC) (reviewed in ^1^). PC is essential for long-range electron transfer ^2–5^, thereby connecting *b_6_f* and PSI during LEF and ensuring that proton motive force and NADPH formation are balanced. This is important, as efficient carbon assimilation requires a fine balance between ATP and NADPH supply, achieved by adjusting the partitioning between LEF and cyclic electron flow (CEF). While LEF produces both ATP and NADPH, CEF exclusively generates ATP, thereby rebalancing the cellular energy budget and contributing to photoprotection through ΔpH regulation ^6–8^.

The so-called “photosynthetic control” arises from lumen acidification, which slows PQH_2_ oxidation at the Qo site of *b_6_f* under low pH conditions ^9,10^. This mechanism modulates electron flow, contributes to non-photochemical quenching ^11–13^, and plays a central role in photoprotection. State transitions provide an additional level of dynamic regulation, mediated in *C. reinhardtii* by the thylakoid-bound STT7 kinase. The thiol-modulated ^14^ and feedback-regulated ^15^ kinase physically interacts with *b_6_f* ^16,17^ to be activated under reduced PQ pool conditions ^18,19^. Activated STT7 phosphorylates several thylakoid membrane proteins, including LHCII, thereby promoting optimized energy distribution between PSII and PSI.

Within chloroplasts of the green alga *Chlamydomonas reinhardtii*, thylakoid membranes are organized into appressed and non-appressed regions, giving rise to lateral heterogeneity in the distribution of photosynthetic complexes ^20,21^. PSII and its LHCII antenna are largely confined to the appressed membranes, whereas PSI and ATP synthase are restricted to non-appressed regions. The *b_6_f* is found in both domains, although its relative abundance has been observed to shift in response to environmental and metabolic cues ^21,22^. Variable accessibility of the stromal gap between appressed thylakoids has been proposed to alter *b_6_f* localization ^23^. In both *C. reinhardtii* and plant chloroplasts adapted to light, the width of the stromal gap is 3.2 – 3.6 nm ^3,24,25^. The *b_6_f* structure has small stromal protrusions (made from the termini and loops of subunit-IV, cytochrome-*b*_6_, and cytochrome-*f*, also known as PetD, PetB, and PetA, respectively ^26^) that extend approximately 1 nm into the stromal gap. We hypothesized that adding a bulky tag to this stromal protrusion might sterically restrict access of *b_6_f* to the stromal gap, thereby excluding it from the appressed thylakoid regions.

In this study, we investigated how modification of the stromal domain of *b_6_f* – by fusing the C-terminal region of PetA to the fluorescent protein (FP) Clover or ATeam – affects (i) photosynthetic electron transfer capacity, (ii) the function of the STT7 kinase to regulate state transitions, and (iii) the spatial organization of *b_6_f* within native thylakoid membranes.

## Results

Stroma-exposed Clover (Extended data Fig. 1a), a bright and photostable FP ^27^, or ATeam (*Adenosine Triphosphate Energy Assay Monitor*) ^28^, a fluorescent sensor of Mg-ATP^2-^, were genetically fused to the C-terminus of PetA (Fig. 1a). Confocal microscopy revealed that, in contrast to WT, ATeam-related fluorescence emission was detectable in independent transformants PetA-ATeam 1 and 2 (Extended data Fig. 2a). Likewise, Clover dependent fluorescence emission was detected in transformants PetA-Clover 1 and 2 (Extended data Fig. 2b). The FP emission co-localized with the chlorophyll fluorescence in the engineered strains, suggesting that the PetA-FP fusions were embedded in the thylakoid membrane as expected. SDS-PAGE followed by immunoblotting confirmed the PetA fusions, where wild-type PetA (36 kDa) was replaced by PetA-ATeam (∼105 kDa) or PetA-Clover (∼65 kDa) (Extended data Fig. 3a-b). Spot test on TP plates, requiring photoautotrophic growth, revealed that PetA-ATeam as well as PetA-Clover transformants performed as WT under normal and high light conditions of 40 and 500 µmol photons m^-2^ s^-1^, respectively (Extended data Fig. 3c-d).

**Fig. 1.**
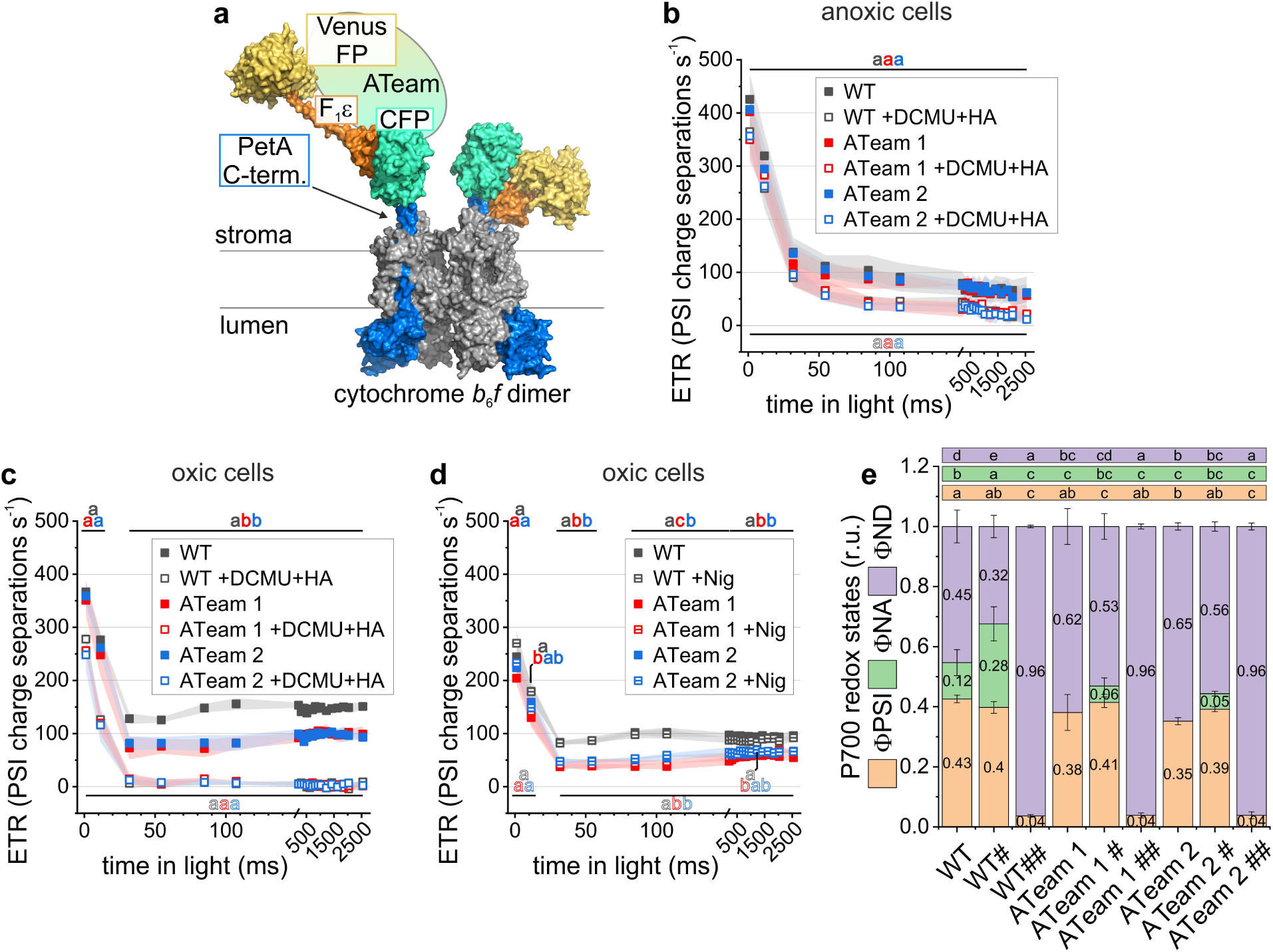
Photosynthetic activities of the cytochrome *b*_6_*f* ATeam variants. (**a**) Structural model, in surface representation, showing the fusion site of ATeam1.03-nD/nA to the PetA of *b*_6_*f* (PDB: 1Q90; PetA in blue). ATeam consists of cyan (CFP; PDB: 2WSN) and yellow fluorescent proteins (Venus; PDB: 3EKJ) joined by the s-subunits of *B. subtilis* ATP synthase. The latter assumes a free conformation (PDB: 4XD7; left) and a MgATP^2-^-bound one (PDB: 2E5Y; right). The functional photosynthesis measurements depict averaged kinetics (*N* = 3 independent biological replicates; mean ± SD; two independent transformants). Electron transfer rates (ETR) developed over the 2.5-s illumination period under (**b**) anoxic and (**c**) oxic conditions in the absence and presence of active PSII. The latter was inhibited by hydroxyl amine (HA) and 3-(3,4-Dichlorophenyl)-1,1-dimethylurea (DCMU). Different letters indicate statistically significant differences among transient ETRs (*N* = 3 independent biological replicates; mean ± SD; one-way ANOVA with Tukey’s HSD post hoc test, *P* < 0.05). Filled and open letters denote conditions without and with added chemicals, respectively. (**d**) ETR were measured under oxic conditions in the absence and presence of 10 µM nigericin (Nig), acting as H^+^/K^+^ exchanger to attenuate ΔpH in favor of an electric field. (**e**) Redox measurements of primary PSI donor, P700, were calculated under conditions shown in panel c. Where indicated, # contained nigericin, and ## signifies additional PSII inhibition by HA and DCMU. The PSI quantum yield (ФPSI) as well as redox-inactive P700 due to acceptor-(ФNA) and donor-side limitation (ФND) are statistically evaluated (One-Way ANOVA/Tukey-HSD, *P* < 0.05).

Next, we measured the electron transfer rate (ETR) in photoautotrophic cells via electrochromic shift (ECS) signals. Under anaerobic conditions that promote transition to State 2, high initial ETR were detected in all strains before stabilizing at a lower steady-state level (Fig. 1b and Extended data Fig. 1b). The ETR toward the end of the measurements ranged from ∼55 to ∼75 PSI charge separations s^-1^, yielding very similar values in WT and engineered PetA fusion strains. Steady-state ETR was substantially decreased when PSII was inhibited, again to a similar level in WT, PetA-ATeam (Fig. 1b), and PetA-Clover strains (Extended data Fig. 1b). The situation differed under oxic conditions, favouring State 1. Approximately 30-50 ms after light exposure, a significant ETR slowdown was detected in PetA-ATeam (Fig. 1c) and PetA-Clover transformants (Extended data Fig. 1c). The steady-state ETR in the fusion strains were ∼60-70% of WT rates. Our attempts to repartition ΔpH for ΔT by treatment with the H^+^/K^+^ exchanger nigericin did not alter this difference (Fig. 1d and Extended data Fig. 1d), suggesting that photosynthetic control was likely not responsible for the ETR slowdown in the engineered PetA fusion strains. We also monitored redox kinetics of the primary PSI donor, P700, under oxic conditions in the presence and absence of nigericin (Fig. 1e, Extended Data Fig. 1e). The data show that P700 donor-side limitation was significantly more pronounced in PetA-ATeam and PetA-Clover transformants than in WT. Additionally, donor-side limitation was less effectively diminished after nigericin treatment compared to WT. Moreover, we measured substantially reduced PSII quantum efficiencies in PetA-ATeam and PetA-Clover transformants compared to WT (Extendend Data Fig. 4). The observed shortcomings in the PetA fusion strains affect the whole photosynthetic machinery under oxic conditions, i.e., lower PSII efficiency and ETR, as well as the increased PSI donor-side limitation that was partially resistant to nigericin. This could be linked to a potential impact on *b_6_f* electron transfer and/or changes in *b_6_f* localization, which could negatively influence PQH_2_ and/or PC-dependent electron transfer between PSII in appressed and PSI in non-appressed thylakoid membrane regions. To determine whether electron transfer within *b_6_f* was affected, we monitored *b_6_f* electrogenicity after single-turnover flashes (Extended Data Fig. 5). The *b_6_f*-dependent ECS rise in WT and both transformants was very similar, indicating that *b_6_f* function remained intact in the engineered strains. Together, the spectroscopic measurements argue against impaired *b_6_f* function in the FP fusion strains as the cause for the electron-transfer dysregulation.

To further explore consequences of the FP fusions, we performed state transition experiments. 77 K fluorescence emission spectroscopy showed an increased PSI antenna size under State 2 versus State 1 conditions in WT cells, but not in PetA-ATeam (Fig. 2a-b) or PetA-Clover strains (Extended Data Fig. 6a-b). This suggests impaired state transitions due to defective STT7-dependent phosphorylation. Indeed, phospho-proteomics in the PetA fusion strains confirmed compromised STT7-dependent phosphorylation of photosystem and LHC targets (see Extended Data Proteomics Results and Extended Data Fig. 7, Extended Data Tab. 1). To check whether this contributed to the photosynthetic phenotypes in Fig. 1, we created *stt7-17*:PetA-ATeam double mutants (Extended Data Fig. 6c). We note that deletion of STT7 in the presence of PetA-ATeam did not prevent rapid P700 oxidation at the onset of illumination, manifested as transient donor side limitation (Fig. 2c). P700 redox kinetics were identical when PSII was inhibited (Extended Data Fig. 6d). Moreover, PetA-ATeam fusion caused a drop in ETR to 60-70% compared to *stt7-17* (Fig. 2d). The same impairment in ETR were observed in PetA-ATeam strains engineered into the CC-125 reference (panels c and e in Fig. 2). These findings rule out altered STT7-dependent protein phosphorylation linked to state transitions as the cause of the reduced LEF in the PetA-FP fusion strains.

**Fig. 2.**
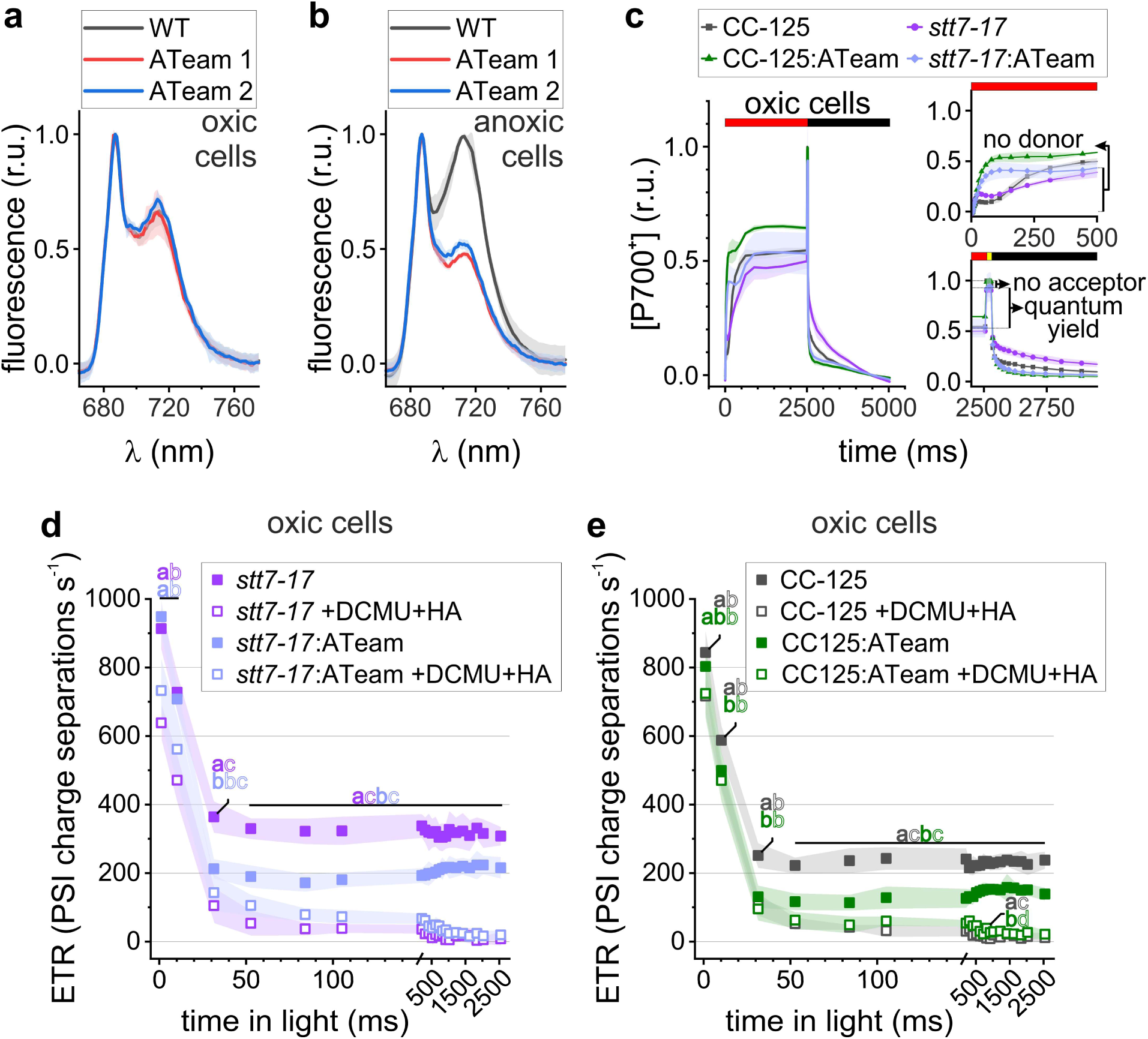
State transition measurements in PetA-ATeam fusion strains and photosynthetic measurements of double mutants lacking STT7 kinase. (**a**, **b**) Differences in antenna attached to photosystem (PS) I under State 1 (oxic) and State 2 (anoxic), contributing to 77 K chlorophyll fluorescence emission at ∼712 nm. These spectra were averaged and normalized to PSII-attached antenna emission signals at ∼687 nm (biological replicates *N* = 3 ± SD). (**c**) Averaged P700 redox kinetics in PetA-ATeam fusion strains obtained in *stt7-17* and its reference CC-125 (*N* = 3 independent biological replicates, each measured in duplicate; mean ± SD). Actinic light, saturating pulse, and darkness are indicated via red, yellow, and black bars, respectively. For the double mutant kinetics, the amplitudes of PSI donor-and acceptor-side limitation, as well as the quantum yield, are labelled. (**d**, **e**) Effect of PetA-ATeam fusion on the electron transfer rate (ETR) time course during 2.5 s of illumination under oxic conditions, in the absence or presence of STT7. Where indicated, PSII was inhibited by hydroxyl amine (HA) and 3-(3,4-Dichlorophenyl)-1,1-dimethylurea (DCMU). Different letters indicate statistically significant differences among transient ETRs (*N* = 3 independent biological replicates, each measured in duplicate; mean ± SD; one-way ANOVA with Tukey’s HSD post hoc test, *P* < 0.05). Filled and open letters denote conditions without and with added chemicals, respectively.

In contrast to the phosphorylation-related changes previously associated with STT7 deficiency ^15^, whole-proteome analysis revealed a pronounced accumulation of proteins involved in central carbon metabolism in the PetA-ATeam strain. These included enzymes of the Calvin– Benson cycle (GAP3, FBA3, FBP2, SEBP1, RPI1, PRK1, PGK1, and TRXf2), starch biosynthesis (PGM1, STA1, and STA6), and carbon mobilization (CAH3, PGL2, and SQD1). Most of these proteins were present at levels approximately 1.5–2-fold higher than in WT cells, whereas ANR1 was ∼3-fold more abundant in WT. In contrast, the PetA-Clover strain showed only minor changes in carbon metabolism proteins, although ANR1 abundance was reduced and NDA3 levels increased by more than 50%. Although annotated as mitochondrial, NDA3 likely localizes to the chloroplast ^29,30^ (see Extended Data Proteomics Results and Extended Data Fig. 7, Extended Data Tab. 1).

To determine whether the PetA-ATeam fusion alters the spatial organization of *b_6_f* within thylakoid membranes, we analyzed appressed membrane regions by cryo-electron tomography (cryo-ET) in WT and PetA-ATeam cells. Particles corresponding to PSII, *b_6_f*, or unassigned complexes were identified by visualization of densities protruding from segmented membrane surfaces (i.e., “membranograms”), followed by manual picking and classification from technical and biological replicates, yielding over 1,000 particles per strain (see Materials and Methods). Representative tomograms, “membranograms”, and assigned particles are shown in (Fig. 3a, 3d, 3e, Extended Data Fig. 8). In WT appressed membranes, PSII and *b_6_f* were present at average concentrations of 777 and 403 particles/μm^2^, respectively, with an additional 318 particles/μm^2^ classified as unassigned, resulting in a total particle density of 1498 particles/μm^2^ (Extended Data Fig. 9). This distribution is consistent with previously reported values ^21,22^, with minor differences likely reflecting strain background or growth conditions. In PetA-ATeam membranes, the total particle density was unchanged (1464 particles/μm^2^), indicating preserved overall membrane protein packing in the appressed regions. However, the relative composition of complexes was significantly altered: PSII concentration increased 28% to 994 particles/μm^2^, while *b_6_f* concentration was reduced by approximately 50% to 193 particles/μm^2^ (Extended Data Fig. 9b). Importantly, the fraction of unassigned particles remained unchanged between strains.

**Fig. 3.**
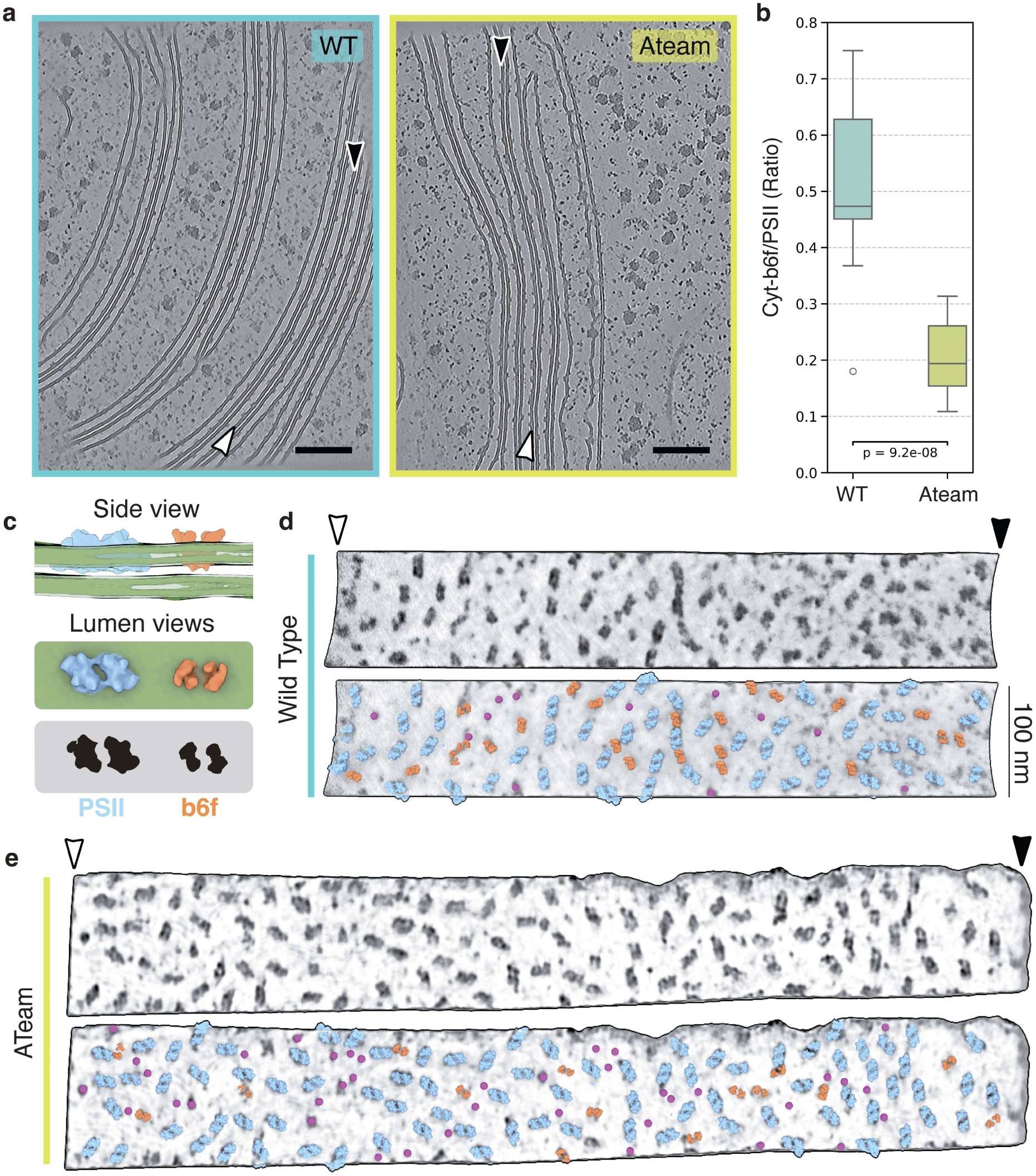
Redistribution of cytochrome *b_6_f* in appressed thylakoid membranes revealed by cryo-electron tomography. (**a**) Representative *XY* slices through cryo-ET volumes from wild-type (WT) and PetA-ATeam cells, showing multiple appressed thylakoid membrane stacks. For each condition, the appressed membrane indicated by a white and a black arrowhead (marking two extremities of the same membrane segment) corresponds to the “membranogram” in panels D and E. Scale bar, 200 nm. (**b**) Ratio of *b_6_f* to PSII particle concentrations measured in appressed thylakoid membranes from WT and PetA-ATeam cells. Boxes indicate the interquartile range, center lines the median, and whiskers the range. Statistical significance was assessed by a two-sided test (p = 9.2 × 10⁻⁸). (**c**) Schematic illustration of PSII and *b_6_f* organization within appressed thylakoid membranes. Top, side view indicating the relative position of the complexes within the membrane stack. Middle, lumen-facing views of PSII (blue) and *b_6_f* (orange). Bottom, corresponding projected “footprints” used for particle identification after intensity projection of tomographic voxels onto segmented membrane surfaces. (**d**) Representative WT appressed thylakoid membrane segment highlighted in panel A. Top, “membranogram” intensity projection of tomogram density onto the segmented membrane surface, revealing proteins protruding from the membrane into the thylakoid lumen. Bottom, the same membrane segment with manually assigned particles overlaid: PSII (blue), *b_6_f* (orange), and unassigned particles (magenta). Scale bar, 100 nm. (**e**) Representative PetA-ATeam appressed thylakoid membrane segment highlighted in panel A, displayed as in panel D. Top, “membranogram” tomogram projection. Bottom, manually assigned PSII (blue), *b_6_f* (orange), and unassigned particles (magenta). Scale bar, 100 nm.

## Discussion

Our cryo-ET data revealed that the increase in molecular mass caused by genetic tethering of ATeam to *b_6_f* on the stromal side altered the distribution of *b_6_f* between appressed and non-appressed thylakoid membranes. These changes in molecular organization are consistent with diminished total ETR, impaired PSII efficiency, and increased donor-side limitations of PSI.

These functional impairments were not seen under strongly reducing conditions such as anoxia. The impact on LEF was not attributable to changes in overall photosynthetic control or *b_6_f* functionality (Fig. 1, Extended data Figs. 1 and 5), nor was it linked to an overall depletion of the tagged *b_6_f* (Extended data Fig. 7). Although the fluorescent PetA-fusion strains failed to undergo state transitions, impairment of STT7-mediated phosphorylation was not the cause of deregulated photosynthesis (Fig. 2 and Extended data Figs. 6 and 7). Given that STT7 associates with the stromal side of *b_6_f*, the modified PetA C-terminus or the enlarged stromal domain of the fusion proteins may sterically hinder complex formation or interfere with conformational changes essential for kinase activation and function. The enrichment of TRXf2 together with Calvin– Benson enzymes is particularly intriguing. It has been suggested that CrTRXf2 is very efficient at targeting Calvin–Benson enzymes for reduction such as FBPase, GAPDH, SBPase, PRK, and PGK ^31^. Increased abundance of both TRXf2 and its targets suggests that the chloroplast may be attempting to maintain reductive activation of the Calvin cycle, because of impaired ETR.

Importantly, the impact of the FP fusions on LEF were likely linked to an altered distribution of *b_6_f* between appressed and non-appressed thylakoid membranes. Cryo-ET analysis of native *C. reinhardtii* cells showed that, although the overall concentration of protein complexes within appressed membranes was similar between WT and PetA-ATeam strains, the relative distribution of PSII and *b_6_f* complexes was markedly shifted. In PetA-ATeam, PSII density in appressed domains increased modestly, while *b_6_f* was substantially depleted, resulting in a marked shift in the PSII/*b_6_f* ratio—from ∼0.52 in WT to ∼0.20 in PetA-ATeam (Figure 3b). This imbalance strongly suggests that the bulky tags fused to the stromal side of *b_6_f* introduce steric hindrance with the stromal gap, thereby impeding their access to the appressed regions. As a consequence, less *b_6_f* is available for LET in the appressed membranes, both to accept PQH_2_ derived electrons from PSII and to donate electrons to PC, which is essential for long-range electron transfer to PSI in non-appressed membranes ^2–4^. The constantly increased PSI donor-side limitation observed in PetA-ATeam and PetA-Clover strains (Fig. 1e, Extended Data Fig. 1e) indicates an electron shortage in the high-potential chain. One plausible cause is non-productive PC exchange between appressed and non-appressed thylakoid regions, which would restrict PC reoxidation by PSI. A non-exclusive alternative is impaired PQH_2_ oxidation at *b_6_f* due to long-range diffusion in the membrane, consistent with the reduced PSII quantum efficiency (Extended Data Fig. 4). Together, the data suggest that restricted PQH_2_ oxidation and/or PC reduction within appressed membranes, potentially due to lower local *b_6_f* concentration in close proximity to PSII, limits electron flow through the chain and ultimately slows LEF (Fig. 4).

**Fig. 4.**
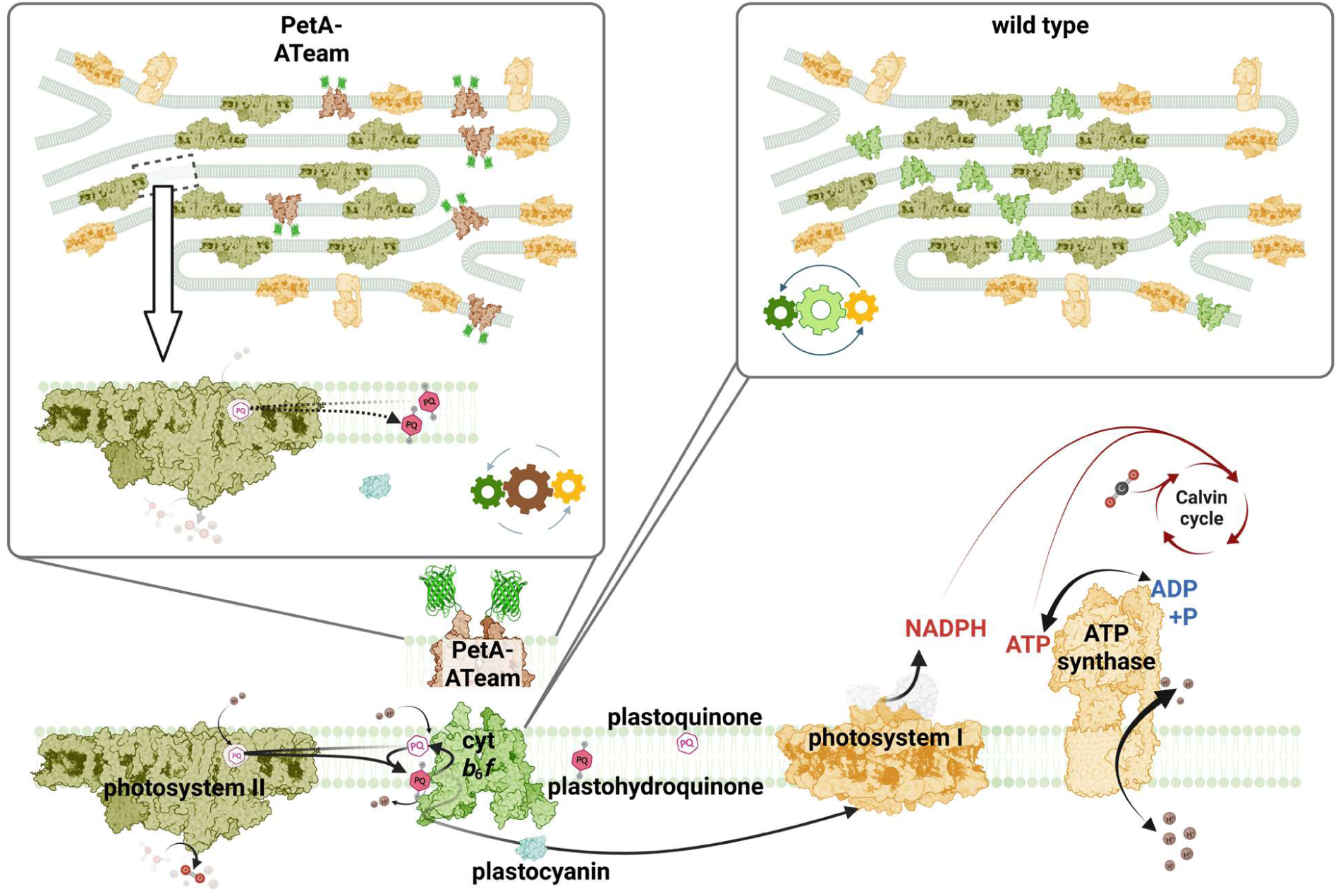
Ultrastructural and functional consequences of artificial redistribution of cytochrome *b_6_f* complex. Depletion of *b*_6_*f* from appressed membrane regions is triggered upon bulky fluorescent protein fusions to the stromal *b*_6_*f* domain. This causes diffusion-limited redox imbalance of mobile electron carriers plastoquinone and/or plastocyanin.

It is important to note that our cryo-ET analysis is restricted to the molecular organization of individual appressed regions, and we cannot rule out broader alterations in the ratio of appressed to non-appressed membrane domains in PetA-ATeam. A comprehensive membrane-wide mapping approach would be required to address this possibility, but this is not feasible by cryo-ET because of the technical requirement to image samples that have been thinned to about <200 nm.

Together, these findings demonstrate that lateral heterogeneity of *b_6_f* in thylakoid membranes is critical for maintaining efficient photosynthetic electron transfer and regulatory flexibility in *Chlamydomonas reinhardtii.* They further indicate that even minor steric perturbations that mislocalize a single electron-transfer complex can substantially impair photosynthetic performance and redox regulation, underscoring membrane organization as a key structural determinant of photosynthetic bioenergetic regulation.

## Methods

### Plasmid Construction

A ∼1.68-kb plastid DNA fragment encompassing the modified petA region of the H6F5 strain ^32^ was amplified using primers petA-5′-F and petA-XbaI-R, with the latter introducing an XbaI restriction site (see Extended Data Tab. 1). The amplicon, whose 3′ *petA* region contained six histidine codons followed by the *petA* stop codon and an NheI site ^32^, was cloned into SmaI/XbaI-digested pUC18 to generate pUC18-petA-H6F5. To replace the six histidine codons with an XhoI site, a 267-bp fragment was amplified from pUC18-petA-H6F5 using primers petA-internal-F and petA-XhoI-R. The PCR product and pUC18-petA-H6F5 were digested with NheI and BsrGI and ligated to generate pUC18-petA-XhoI-NheI. Insertion of the spectinomycin-resistance marker was performed through an intermediate construct derived from pUC18-aadA ^15^. An 881-bp fragment extending downstream of *petA* was amplified using primers petA-3′-F and petA-3′-R and blunt-end ligated into SmaI-digested pUC18-aadA, yielding pUC18-aadA-881. A 2.38-kb fragment comprising the aadA cassette and the downstream *petA* region was subsequently amplified from this construct using primers petA-3′-R and aadA-XbaI-F, with the latter introducing an XbaI site. The resulting fragment was digested with XbaI, whereas pUC18-petA-XhoI-NheI was linearized with SbfI, blunt-ended using T4 DNA polymerase, and subsequently digested with XbaI. Ligation of the two fragments generated pUC18-petA-aadA, containing XhoI and NheI sites for the introduction of *petA* gene fusions. The linker peptide for fusion proteins was SSSAGSAAGSGEF. Starting codons of Clover (GenBank: AFR60231.1) and ATeam1.03-nD/nA (Addgene plasmid # 51958) were omitted. Codon optimization was carried out with CSO software ^33^ and synthetic genes plus the linker sequence (Thermo Fisher Scientific) were directly cloned into XhoI/NheI-digested vector to yield pUC18-petA-Clover-aadA and pUC18-petA-ATeam-aadA, respectively (for synthesized DNA sequences, see Extended Data Tab. 1).

### Biolistic Transformation of WT Strain

Besides CC-124 (mt -), *stt7-17* ^15^ and its reference CC-125 (mt +) served as recipients. Mutant strains were generated through chloroplast transformation using plasmids containing the modified *petA* gene version along with an aadA cassette to confer spectinomycin resistance (150 µg/ml). Spectinomycin resistant clones were selected and replated several times to obtain homoplastic strains using primers petA-internal-F and petA-XbaI-R (see Extended Data Tab. 1).

### Growth conditions and treatments

Cells were maintained at 50 µmol photons m^-2^ s^-1^ at 23°C on agar plates of Tris-acetate-phosphate (TAP) medium or Tris-minimal medium (TP) in the absence of acetate ^34^ by bubbling sterile air under a 16-h light/8-h dark regime. The effective photosynthetic active radiation was determined with a Quantitherm PAR/Temp Sensor (Hansatech, England). For state transition treatment, TAP-grown synchronized cultures were freshly diluted one day before the experiment and, upon harvesting (4000 *g*, 5 min, 23°C), adjusted to 5 µg chlorophyll ml^-1^. Aliquots were either kept for 45 min in aerobic conditions by shaking in 40 µmol photons m^−2^ s^−1^ light upon adding 10 µM 3-(3,4-Dichlorophenyl)-1,1-dimethylurea (DCMU; State 1), or were given anoxic treatment in the dark using a glucose oxidase/catalase cocktail (State 2) ^35^. For proteomic analyses, TAP grown cells were shifted to State 2 conditions for 30 min and samples for phospho-proteomic analyses were collected. The strains were checked for photoautotrophic growth for one week by spotting 20 µl of 1×10^6^ cells/ml on TP agar plates at 25°C kept under 40 µmol photons m^−2^ s^−1^ (16 h light/8 h dark) and continuous 500 µmol photons m^−2^ s^−1^. Liquid TP cultures were harvested for optical spectroscopy and Ficol (20% *w*/*v*) was added to the TP resuspension medium.

### Biochemical methods

Liquid cultures of the variant strains were grown in TAP medium and freshly diluted one day before harvesting. The cells were pelleted and mixed in a buffer containing 0.2 M dichloro-diphenyl-trichloroethane, 0.2 M Na_2_CO_3_, 10 mM NaF, and protease inhibitors (0.2 mM phenylmethylsulfonyl fluoride, 1 mM benzamidine, 20 mM amino caproic acid) to be analyzed via SDS-PAGE ^36^. Primary antibodies against PetA (AS06 119) and STT7 (AS15 3080), as well as secondary antibodies (AS09 602) were purchased from Agrisera.

### Sample preparation for mass spectrometry

Protein isolation from cell pellets was carried out as described ^37^. Reduction/alkylation of cysteines and tryptic digestion of proteins (100 µg per sample) was performed following the SP4 protocol ^38^. After digestion, peptides were desalted using PurePep H50 SPE Spin columns (Affinisep). The peptide samples were divided into two aliquots: 5 µg for whole proteome analysis and 95 µg for phosphopeptide enrichment. Both aliquots were dried using vacuum centrifugation before further processing.

To enrich phosphopeptides, titanium dioxide (TiO2) beads with 5 µm particle size (Titansphere, GL Sciences) in a metal oxide affinity chromatography (MOAC) procedure were used. TiO2 (1 mg per sample) was activated once with acetonitrile and equilibrated three times with loading buffer (LB; 80% (v/v) acetonitrile/5% (v/v) TFA/1 M glycolic acid). Loading buffer was added to create a 10% (w/v) TiO2 suspension. Peptide samples were dissolved in 50 µl LB, combined with 10 µl TiO2 suspension, and incubated for 30 minutes at 25°C in an Eppendorf Thermomixer at 1,200 rpm). The mixture was transferred to SDB-XC STAGE tips prepared in-house ^39^. Unless stated otherwise, all subsequent washing and elution steps were performed using 60 µl buffer volumes with in-between centrifugation at 2000 × g at room temperature. The buffer was removed by centrifugation and the TiO2 beads were washed twice with LB and once with 1% (v/v) acetonitrile/0.1% (v/v) TFA (W1). Non-phosphorylated peptides that were removed from the beads and subsequently bound to the SDB-XC membrane plug were removed by one wash with 80% (v/v) acetonitrile/0.1% (v/v) TFA (W2). The phosphopeptides were eluted from the TiO2 beads onto the SDB-XC material using 160 µl of pH 11 buffer (0.4 M sodium hydrogen phosphate/NaOH with 1% acetonitrile). After washing once with 1% acetonitrile/0.1% TFA, the phosphopeptides were sequentially eluted in three fractions using increasing concentrations of acetonitrile (7.5%, 20%, and 60%) in 100 mM ammonium formate (pH10). For samples from the STT7 experiment, only one fraction (60% acetonitrile in 100 mM ammonium formate (pH10)) was collected. The strongly bound phosphopeptides were eluted in two steps: first from the TiO2 beads using 5% ammonia, then from the SDB-CX membrane using 60% acetonitrile in 100 mM ammonium formate at pH 10. All fractions were dried by vacuum centrifugation and stored at −80°C until MS analysis.

### Mass spectrometry

#### Whole proteome analysis

Dried peptide samples were reconstituted in 5 µl of 2% (v/v) acetonitrile/0.05% (v/v) TFA in LC/MS grade water. Samples were analyzed on an LC-MS/MS system consisting of an Ultimate 3000 NanoLC (Thermo Fisher Scientific) coupled via a Nanospray Flex ion source (Thermo Fisher Scientific) to a Q Exactive Plus mass spectrometer (Thermo Fisher Scientific). Peptides were concentrated on a trap column (Acclaim Pepmap C18, 5 × 0.3 mm, 3 µm particle size, Thermo Scientific) for 3 min using 4% (v/v) acetonitrile/0.05% (v/v) in LC/MS grade water at a flow rate of 10 µl/min. The trap column was operated in back-flush mode, allowing the transfer of peptides on a reversed-phase column (PepSep Fifty, 500 × 0.075 mm, 1.9 µm particle size, Bruker) for chromatographic separation. The eluents used were 0.1% (v/v) formic acid in LC/MS grade water (A) and 0.1% (v/v) formic acid/80% (v/v) acetonitrile in LC/MS grade water (B). The gradient was programmed as follows: 2.5% to 5% B over 5 min, 5% to 17.5% B over 47 min, 17.5% to 40% B over 105 min, 40% to 99% B over 10 min, and 99% B for 20 min. Flow rate was 250 nl/min. The mass spectrometer was operated in data-dependent acquisition (DDA) mod, alternating between one MS1 full scan and up to 12 MS2 scans. Full scans were acquired with the following settings: AGC target 3e6, MS1 resolution 70000, maximum injection time 50 ms, scan range: 350–1400 m/z. Settings for MS were: AGC target 5e4, resolution 17500, MS2 maximum injection time 80 ms, intensity threshold 1e4, normalized collision energy 27 (HCD)). Dynamic exclusion was set to ‘auto’, assuming a chromatographic peak width (FWHM) of 60 s.

### Phosphoproteomic analysis

Prior to analysis, dried peptide samples were reconstituted in 5 µl of 2% (v/v) acetonitrile/0.05% (v/v) TFA in LC/MS grade water. LC-MS/MS system, flow rate and eluent compositions were the same as described above. The gradient for peptide separation was as follows: 2.5% to 5% B over 5 min, 5% to 40% B over 92 min, 40% to 99% B over 10 min, and 99% B for 20 min. The mass spectrometer used the same DDA mode settings as the whole proteome analysis, with two changes: MS2 maximum injection time was increased to 120 ms, and dynamic exclusion was set to 45 s.

### MS data analysis

For quantitative phosphoproteome analysis, mass spectrometry raw data from nonenriched and enriched samples were searched using Fragpipe ^40,41^ against a concatenated, non-redundant database containing *Chlamydomonas reinhardtii* protein sequences from v5.6 and v6.1 gene models (Phytozome 13, phytozome-next.jgi.doe.gov). In addition, the polypeptide sequences of mutated/tagged proteins were added to the database. Default ‘LFQ-MBR’ workflow settings were applied to data from whole proteome samples, while the ‘LFQ-phospho’ workflow was used for phosphopeptide samples. A false discovery rate of 1% was applied at both peptide and protein levels.

Fragpipe output files containing peptide/protein identifications and quantitative data (’combined_modified_peptide.tsv’ from enriched samples and ‘combined_protein.tsv’ from whole proteome samples) were reformatted using a custom Python script to ensure compatibility with Phospho-Analyst (v. 1.0.2), which was used for differential expression analysis ^42^: Missing values were imputed using the ‘MinProb’ function, and variance stabilizing transformation was applied to normalize phosphosite intensities. Since the whole proteome LFQ data was already normalized in Fragpipe, no additional normalization steps were needed. Phospho-Analyst automatically corrected phosphosite abundance changes for underlying protein levels. A phosphorylation site localization probability threshold of 0.33 was applied, and p-values were adjusted using the Benjamini-Hochberg method. Predicted protein localizations are based on recently published data (PB-Chlamy)^29^.

The mass spectrometry proteomics data have been deposited to the ProteomeXchange Consortium (http://proteomecentral.proteomexchange.org) via the PRIDE partner repository ^43^ with the dataset identifier PXD082283.

### Optical spectroscopy

The measurements were carried out by using a Joliot-type spectrophotometer (JTS-150, SpectroLogiX) equipped with the interference filters and deconvolution procedures as reported previously ^35^. For multiple turnover conditions, previously described conditions and protocols used cells which were adapted to alternating 2.5 s actinic light (AL; 550 µmol photons m^-2^ s^-1^ peaking at 630 nm) and dark. Briefly, ETR were obtained in the presence of LEF but referred to PSI charge separation activity. This stems from separate ECS calibration measurements that allowed, via saturating single-turnover flashes (Q-switched Nd:YAG, Continuum), for the conversion of optical changes (obtained as Δ*I*/*I* at 520 nm −546nm) into the ECS signals corresponding to 1 PSI charge separation ^44^. Normalization routines were carried out in the presence of PSII inhibitors hydroxyl amine (HA; 2 mM) and 3-(3,4-Dichlorophenyl)-1,1-dimethylurea (DCMU; 20 µM). The ETR protocol was based on a Dark Interval Relaxation Kinetics approach ^45,46^. P700 measurements during 2.5 s AL and dark relied on the Klughammer & Schreiber method ^47^.

### Low-temperature chlorophyll fluorescence measurements

77 K fluorescence emission spectra of whole cells were obtained in liquid nitrogen in a FP-6500 spectrofluorometer (Jasco, Germany).

### Sample preparation for cryo electron tomography (cryo-ET) and plunge-freezing

*Chlamydomonas reinhardtii* cells, either wild-type (CC-124) or ATeam strain expressing a genomically integrated PetA-Ateam tag (CC-124 background), were grown independently in two biological replicates. Each replicate was cultured in Tris-acetate-phosphate (TAP) medium for 2– 4 days, then diluted 1:20 into minimal medium and incubated under continuous illumination (60 μmol photons m⁻^2^ s⁻¹) with vigorous shaking. For each replicate, 4.2 μl aliquots of log-phase cells were applied to glow-discharged Quantifoil R2/1 carbon-coated copper 200 mesh grids (Quantifoil Micro Tools). Plunge-freezing was performed on three separate occasions using a Vitrobot Mark IV (Thermo Fisher Scientific) set to 22°C and 95% humidity, with a blot time of 7 s. Grids were vitrified in a liquid ethane and clipped into AutoGrids (Thermo Fisher Scientific) for cryo-FIB milling.

### Focused Ion Beam (FIB) milling

Cryo-lamellae were prepared using an Aquilos 2 dual-beam FIB/SEM instrument (Thermo Fisher Scientific) ^48^. Grids were first coated with a thin protective layer of organometallic platinum using the integrated gas injection system (GIS). Lamellae were milled with a gallium ion beam using a stepwise current reduction from 1.5 nA to 50 pA, targeting a final lamella thickness of 120–180 nm.

### Cryo-ET data acquisition

Tilt-series were acquired on two Titan Krios 300 kV transmission electron microscopes (Thermo Fisher Scientific), both equipped with a Falcon 4 direct electron detector and a Selectris X energy filter operated with a 10 eV slit. One of the instruments was additionally equipped with a monochromator. Data sets were collected using TEM Tomography 5 software (Thermo Fisher Scientific) in EER format with a dose-symmetric tilt scheme ^49^, with a pretilt of ±10° and 2° increments and a total span up to 120°. Images were recorded at a nominal pixel size of 1.95 Å or 1.98 Å. The cumulative electron dose per tilt-series was maintained between 120 and 140 e⁻/Å^2^, and target defocus values of tilt series ranged from −2.5 μm to −5 μm. To ensure biological and technical independence, tomograms were collected over four distinct sessions and each tomogram was acquired on a separately milled cell.

### Tomogram reconstruction and membrane segmentation

Raw movie frames were motion-corrected using MotionCor2 ^50^. Dose-weighting and contrast transfer function (CTF) estimation were performed in IMOD (v4.11) ^51^. Fiducial-less tilt-series alignment was carried out using AreTomo ^52^. Tomograms were reconstructed by weighted back-projection using IMOD ^53^ at a final pixel size of 7.8 Å or 7.92 Å (bin4). For contrast enhancement and partial recovery of missing wedge information, DeepDeWedge ^54^ was applied. The neural network was trained and applied to each tomogram individually, using extracted sub-tomograms of 80 × 80 × 80 voxels.

To isolate biologically-relevant tomogram volume, binary masks encompassing only the lamella were generated using Slabify ^55^, and these were used to multiply the reconstructed tomograms. The masked tomograms were then processed using the MemBrain-seg module from MemBrain v2 ^56^, to segment all thylakoid membranes. For downstream structural analysis, individual membrane instances were manually segmented in Amira3D (Thermo Fisher Scientific), exported as binary MRC volumes, and converted into triangulated surface meshes using the *membrain_pick convert_mb_folder* function from MemBrain v2.

### Particle annotation and statistical analysis

For each strain (Chlamydomonas wild-type and ATeam), ∼150 individual membrane segmentations were initially generated, spanning four tomograms (3–4 membranes per tomogram). From these, we selected 16 and 14 appressed membranes from the WT and ATeam datasets, respectively, in which PSII and cytb6f complexes could be annotated with the highest confidence.

A total of 1,194 and 1,233 particles were manually annotated for the WT and ATeam datasets respectively (see Table 2 for details). The annotation was performed on appressed membranes in which PSII and cytb6f complexes could be confidently identified based on structural features and membrane-associated density footprints. Annotation was guided by membrane projections at varying distances from the surface, achieved by dynamically expanding or contracting the surface mesh normal to the membrane using Surforama. Particles were classified into three categories: PSII, cytb6f, or not-assigned (NA). The surface area of each segmented membrane was computed using MemBrain v2 and particle concentrations (particles/μm^2^) was calculated for each membrane individually, using the *membrain_stats* function *–exclude_edges --edge-exclusion-width* set to 50 Å. Particle concentrations were compared between WT and ATeam appressed membranes using violin plots with overlaid swarm plots generated with Python. Statistical significance was assessed using unpaired two-tailed t-tests for each particle class. The ratio of cyt-b6f to PSII particles was calculated per membrane and visualized using box-and-whisker plots. Statistical significance between WT and ATeam was annotated using the corresponding p-value.

### Figure preparation

Statistical plots, including violin, box, and histogram-based visualizations, were generated in Python using Seaborn, Matplotlib, and SciPy, based on exported tables of particle counts and distances. Scripts were customized to annotate significance values and distribution peaks directly onto the plots.

Segmented binary membrane volumes exported from Amira3D were post-processed in IMOD to generate smooth surfaces for rendering. This involved intensity clipping (*clip threshold*) followed by surface smoothing (*clip smooth*). The resulting MRC volumes were opened in ChimeraX ^57^ (v1.8) alongside the corresponding DeepDeWedge-denoised tomograms. Membranes were colored by sampling intensity values from the tomogram using the color sample command with a custom grayscale palette (*palette −1,#000000:-0.3,#333333:0.5,#808080:1,#bfbfbf:3,#e4e4e4*). All membranes were visualized at a consistent isosurface threshold (*surface level 0.153*) across figures. Particle coordinates and orientations were loaded into ArtiaX ^58^, and rendered as volume representations using molmap-generated density models (18 Å resolution) based on PDB entries 6KAC (PSII) and 7QRM (cytb6f). Not-assigned (NA) particles were displayed as red spheres.

## Data availability

Cryo-ET cellular tomograms are available in the Electron Microscopy Data Bank (EMDB) with the accession codes EMD-XXXXX1, EMD-XXXX2, EMD-XXXX3, EMD-XXXX4, EMD-XXXX5, EMD-XXXX6, EMD-XXXX7, and EMD-XXXX8. Raw electron tomography data are available in the Electron Microscopy Public Image Archive (EMPIAR-XXXXX).

## Acknowledgements

M.H. (507704013) and F. B. (507704013) acknowledge funding from the Deutsche Forschungsgemeinschaft (DFG FOR 5573/1). B.D.E. acknowledges funding from the Swiss National Science Foundation (SNSF; project grant 310030E_217510), as a part of DFG FOR 5573/1. D.T. was supported by an EMBO Postdoctoral Fellowship (ALTF 1333-2024) and an SNSF Postdoctoral Fellowship (TMPFP3 233747). M.H. acknowledges the RECTOR program in association with the University of Okayama, Japan.

## Author contributions

M.H., B.D.E. and F.B. designed the project. A.Z. and F.B. carried out genetic engineering, kinetic measurements and analyses. A.Z. and M.Schw. carried out confocal microscopy analyses. M.Scho. did the mass spectrometric measurements and data evaluation. M.Scho. and M.H. analysed the mass spectrometric data. D.T. performed cryo-ET workflow and performed the quantitative analyses. W.W. contributed to cryo-ET grid preparation and tomogram analysis. D.T., W.W., and B.D.E. interpreted the cryo-ET results. F.B. and M.H. provided the initial draft and all authors contributed to the final version of the manuscript.

