## Supplemental Information for "Lateral organization of cytochrome *b_6_f* in thylakoid membranes controls photosynthetic electron transfer efficiency"

**Extended Data Contents**

Extended Data Proteomics Results
Extended Data Proteomics Methods
Extended Data References
Extended Data Figures 1 – 9
Extended Data Table 1 *(separate spreadsheet file)*

**Extended Data Proteomics Results**

Differential expression analysis revealed significantly up- or downregulated phosphorylation sites under State 2 conditions (|$\log_2$ fold change| ≥ 2, FDR < 0.05; Extended Data Fig. 7; Extended Data Table 1). The altered phosphorylation patterns closely resembled those of known *stt7* mutants in *Chlamydomonas reinhardtii* and *Arabidopsis thaliana* [1-4].

Notably, N-terminal phosphorylation of LHCB4 (T7, T11, T7+T11) and LHCSR3 (T32, T33) was strongly reduced in PetA-ATeam strains, reaching only a fraction of WT levels ($\log_2$ fold change 4 to 14.7), with double phosphorylation of LHCSR3 (T32+T33) entirely absent. Other STT7 targets, including PSBR (T37), ALB3.2 (T325), PSAG (T71), PetD (T4), and PETO (T164), showed severely reduced phosphorylation, while PETO (T169) phosphorylation was undetectable. The nucleoid protein pTAC16 displayed nine hypophosphorylated sites in PetA-ATeam. Conversely, phosphorylation of TAP38 (T420), an STT7 antagonist [5], was elevated. PetA-Clover strains exhibited a similar trend with some differences: LHCB4 phosphorylation remained unchanged, single LHCSR3 (T32/T33) phosphorylation was unaffected, but double phosphorylation (T32+T33) was reduced. Unlike PetA-ATeam, pTAC16 showed no significant changes. Interestingly, STT7 was phosphorylated at T147 (or T152) within the ATP-binding site in all PetA-Clover replicates, but not in WT or PetA-ATeam. Five additional STT7 phosphorylation sites (localization > 0.75), located outside the kinase domain, were unchanged across strains. Both fusion proteins carried phosphorylated residues, at least two in PetA-Clover and six in PetA-ATeam (Extended Data Table 1).

Whole-protein analysis revealed few global changes in PetA-ATeam compared to WT, but most affected proteins were central to carbon metabolism, including Calvin–Benson cycle enzymes (GAP3, FBA3, FBP2, SEBP1, RPI1, PRK1, PGK1, TRXf2), starch synthesis (PGM1, STA1, STA6), and carbon mobilization (CAH3, PGL2, SQD1) (Extended Data Fig. 6). These proteins were generally 1.5–2× more abundant in transformants, except ANR1, which was ~3× higher in WT. In PetA-Clover, carbon metabolism proteins were largely unchanged, but ANR1 decreased and NDA3 increased > 50%. Although annotated as mitochondrial, NDA3 likely localizes to the chloroplast [6,7].

### Extended Data References


1 Bergner, S. V. *et al.* STATE TRANSITION7-Dependent Phosphorylation Is Modulated by Changing Environmental Conditions, and Its Absence Triggers Remodeling of Photosynthetic Protein Complexes. *Plant Physiol* **168**, 615-634, doi:10.1104/pp.15.00072 (2015).

2 Lemeille, S. *et al.* Analysis of the chloroplast protein kinase Stt7 during state transitions. *PLoS biology* **7**, e45-e45, doi:10.1371/journal.pbio.1000045 (2009).

3 Lemeille, S., Turkina, M. V., Vener, A. V. & Rochaix, J.-D. Stt7-dependent phosphorylation during state transitions in the green alga Chlamydomonas reinhardtii. *Mol Cell Proteomics* **9**, 1281-1295, doi:10.1074/mcp.M000020-MCP201 (2010).

4 Zaeem, A. *et al.* N-terminal region of PetD is essential for cytochrome b6f function and controls STT7 kinase activity via STT7-dependent feedback loop phosphorylation. *bioRxiv*, doi:doi: https://doi.org/10.1101/2025.02.21.639470 (2025).

5 Cariti, F. *et al.* Regulation of Light Harvesting in Chlamydomonas reinhardtii Two Protein Phosphatases Are Involved in State Transitions. *Plant Physiol* **183**, 1749-1764, doi:10.1104/pp.20.00384 (2020).

6 Wang, L. *et al.* A chloroplast protein atlas reveals punctate structures and spatial organization of biosynthetic pathways. *Cell* **186**, 3499-3518.e3414, doi:https://doi.org/10.1016/j.cell.2023.06.008 (2023).

7 Terashima, M., Specht, M., Naumann, B. & Hippler, M. Characterizing the anaerobic response of Chlamydomonas reinhardtii by quantitative proteomics. *Mol Cell Proteomics* **9**, 1514-1532, doi:10.1074/mcp.M900421-MCP200 (2010).

8 Younas, M., Scholz, M., Marchetti, G. M. & Hippler, M. Remodeling of algal photosystem I through phosphorylation. *Biosci Rep* **43**, doi:10.1042/BSR20220369 (2023).

9 Johnston, H. E. *et al.* Solvent Precipitation SP3 (SP4) Enhances Recovery for Proteomics Sample Preparation without Magnetic Beads. *Anal Chem* **94**, 10320-10328, doi:10.1021/acs.analchem.1c04200 (2022).

10 Rappsilber, J., Mann, M. & Ishihama, Y. Protocol for micro-purification, enrichment, pre-fractionation and storage of peptides for proteomics using StageTips. *Nature Protocols* **2**, 1896-1906, doi:10.1038/nprot.2007.261 (2007).

11 Kong, A. T., Leprevost, F. V., Avtonomov, D. M., Mellacheruvu, D. & Nesvizhskii, A. I. MSFragger: ultrafast and comprehensive peptide identification in mass spectrometry–based proteomics. *Nature Methods* **14**, 513-520, doi:10.1038/nmeth.4256 (2017).

12 Yu, F. *et al.* Analysis of DIA proteomics data using MSFragger-DIA and FragPipe computational platform. *Nat Commun* **14**, 4154, doi:10.1038/s41467-023-39869-5 (2023).

13 Zhang, H. *et al.* Phospho-Analyst: An Interactive, Easy-to-Use Web Platform To Analyze Quantitative Phosphoproteomics Data. *J Proteome Res* **22**, 2890-2899, doi:10.1021/acs.jproteome.3c00186 (2023).

14 Perez-Riverol, Y. *et al.* The PRIDE database and related tools and resources in 2019: improving support for quantification data. *Nucleic acids research* **47**, D442-D450 (2019).

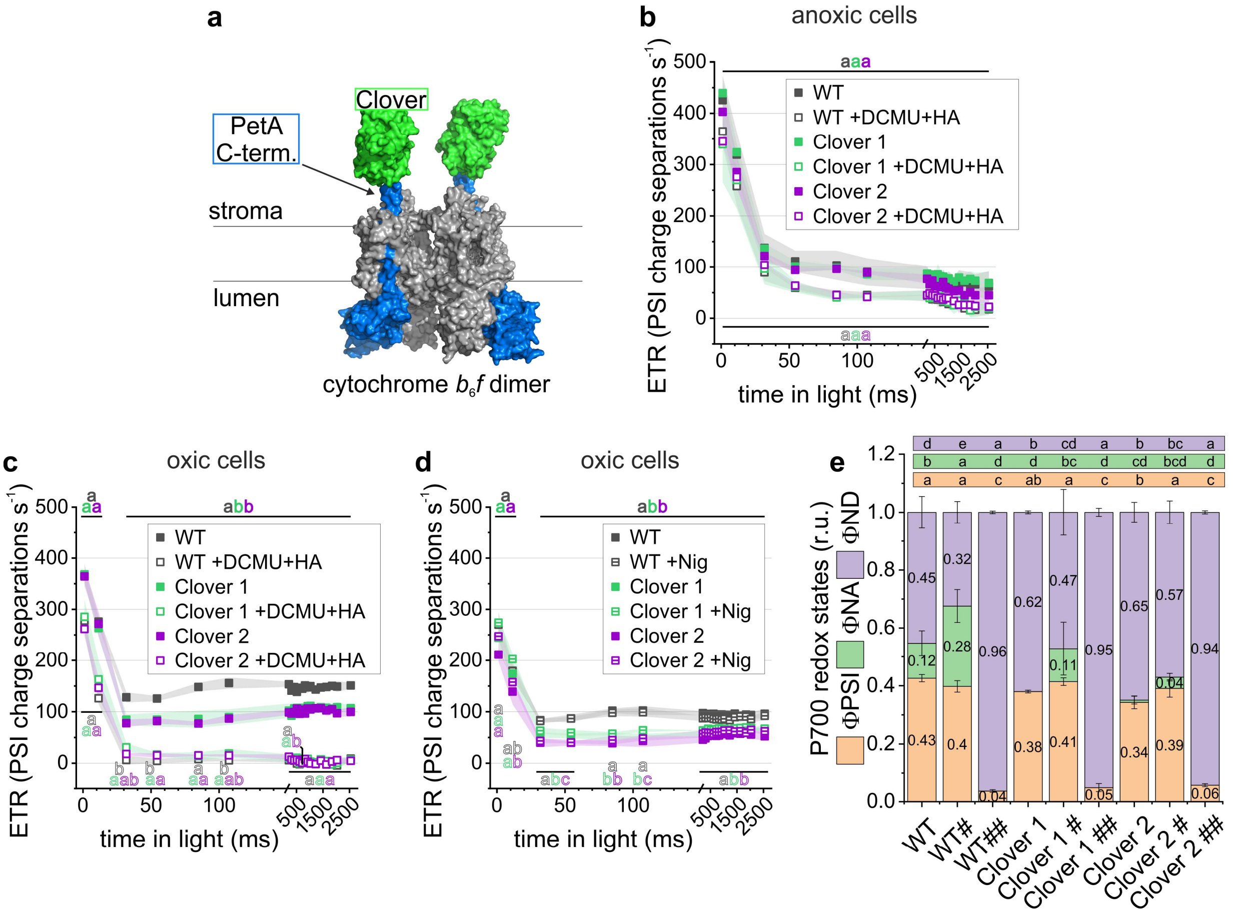


**Extended Data Fig. 1. Photosynthetic activities of the cytochrome $b_6f$ variants containing Clover.** (**a**) Structural model, in surface representation, showing the site of Clover (PDB: 5WJ2) to the PetA of $b_6f$ (PDB: 1Q90; PetA in blue). The functional photosynthesis measurements depict averaged kinetics ($N = 3$ independent biological replicates; mean ± SD; two independent transformants). Electron transfer rates (ETR) developed over the 2.5-s illumination period under (**b**) anoxic and (**a**) oxic conditions in the absence and presence of active PSII. The latter was inhibited by hydroxyl amine (HA) and 3-(3,4-Dichlorophenyl)-1,1-dimethylurea (DCMU). Different letters indicate statistically significant differences among transient ETRs ($N = 3$ independent biological replicates; mean ± SD; one-way ANOVA with Tukey's HSD post hoc test, $P < 0.05$). Filled and open letters denote conditions without and with added chemicals, respectively. (**d**) ETR were measured under oxic conditions in the absence and presence of 10 µM nigericin (Nig), acting as $H^+/K^+$ exchanger to attenuate ΔpH in favor of an electric field. (**e**) Redox measurements of primary PSI donor, P700, were calculated under conditions shown in panel c. Where indicated, # contained nigericin, and ## signifies additional PSII inhibition by HA and DCMU. The PSI quantum yield (ΦPSI) as well as redox-inactive P700 due to acceptor- (ΦNA) and donor-side limitation (ΦND) are statistically evaluated (One-Way ANOVA/Tukey-HSD, $P < 0.05$).

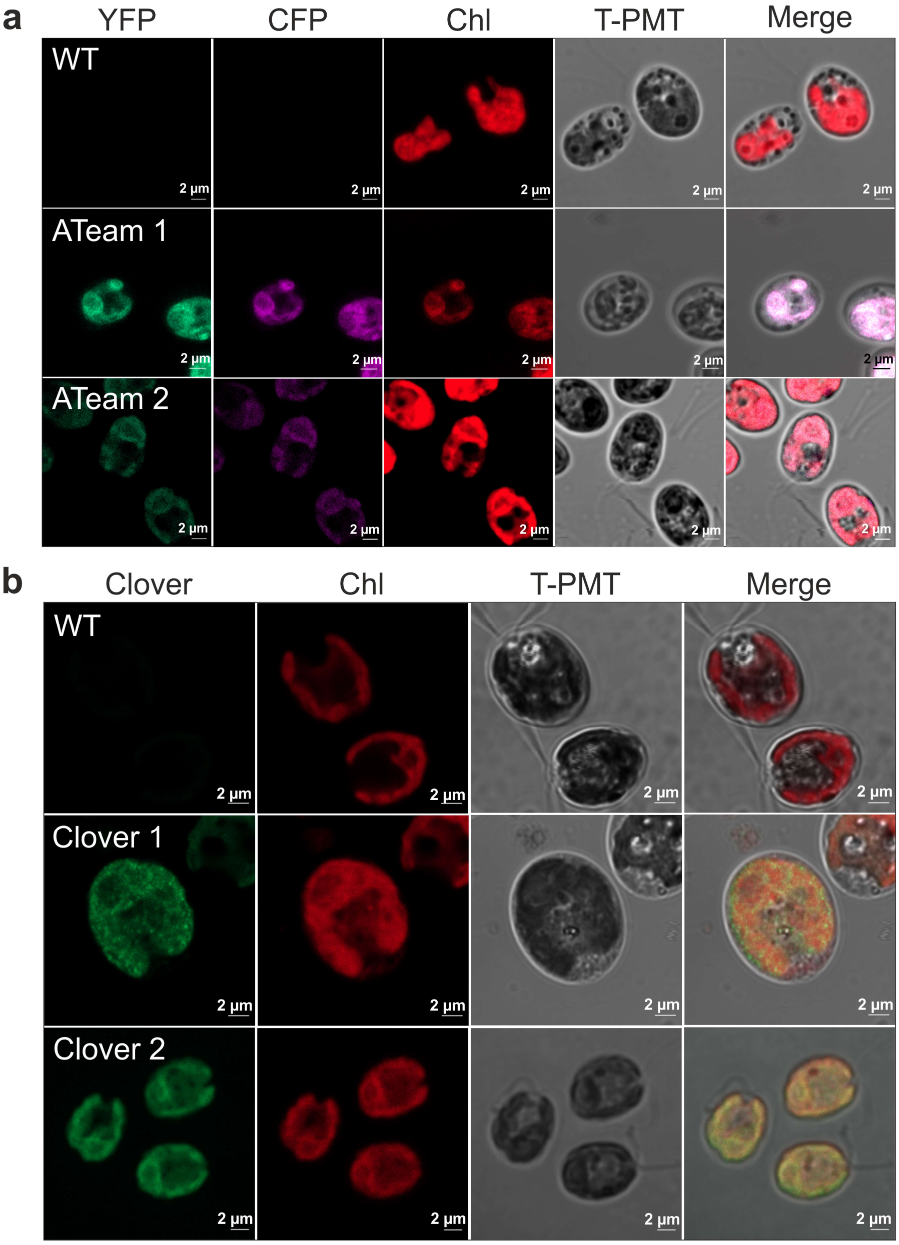


**Extended Data Fig. 2. Confocal microscopy of fluorescent protein expression. (a)** Fluorescence emission from YFP and CFP variants of the ATeam sensor. (**b**) Clover-derived fluorescence in the respective transformants. Chlorophyll auto-fluorescence (Chl) is shown for reference, as well as bright field (T-PMT) and merged channel signals.

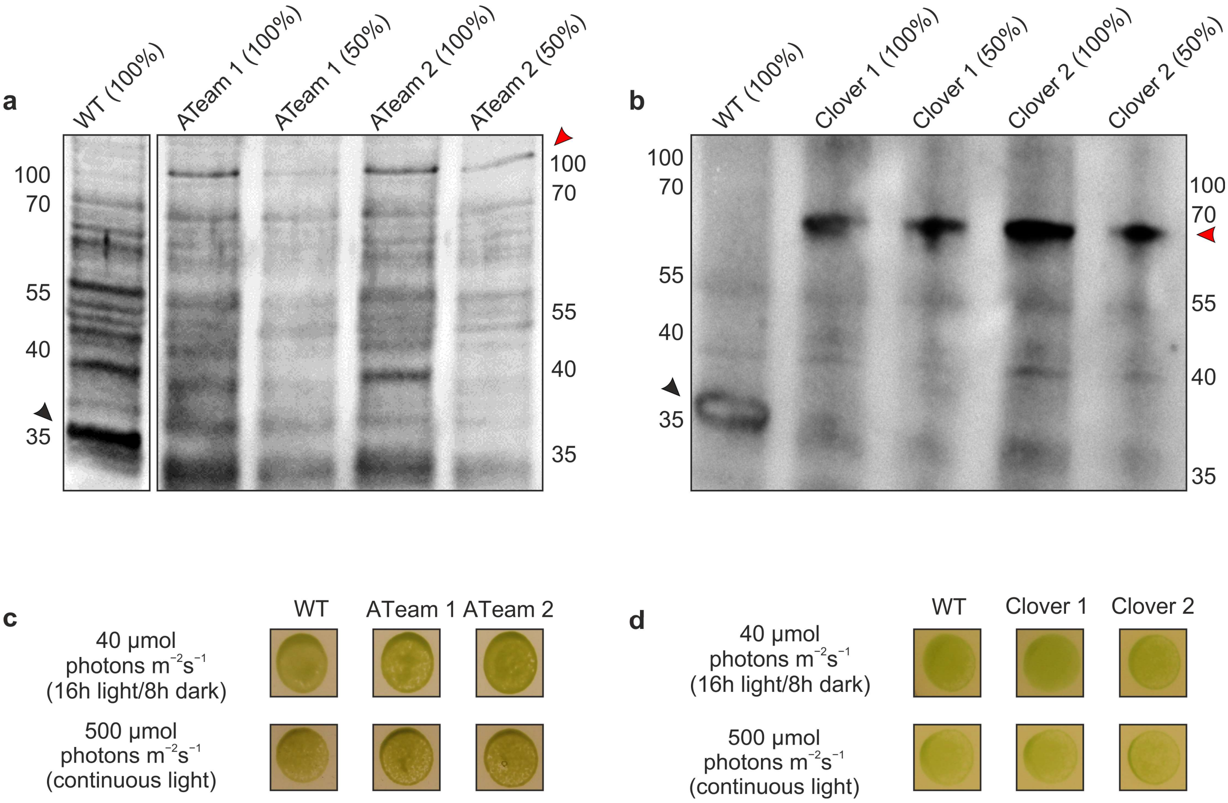


**Extended Data Fig. 3. Immunodetection of PetA as well as growth analysis in varying light conditions with and without GFP fusion.** Successful GFP fusions are visualized via immunodetection of anti-PetA signals for (**a**) ATeam and (**b**) Clover transformants, using whole cell extracts. Spot tests on minimal media compare the growth under two light conditions (40 and 500 µmol photons $m^{-2}$ $s^{-1}$) under diurnal and continuous light regimes, respectively. Representative colonies are shown for (**c**) ATeam and (**d**) Clover transformants.

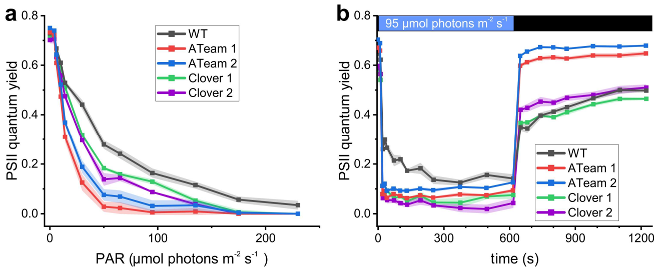


**Extended Data Fig. 4. Chlorophyll fluorescence-based photosynthesis measurements of the cytochrome *b*$_6$*f* variants containing ATeam and Clover.** Averaged kinetics are depicted (biological replicates $N = 3 \pm$ SD; two independent transformants). The PSII quantum yield during (**a**) light curve and (**b**) continuous light induction measurements are shown.

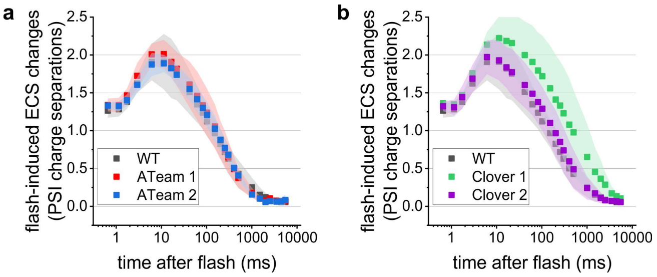


**Extended Data Fig. 5. A rise in electrochromic shift (ECS) measured ~10-ms after a laser flash show $b_6f$-related electrogenicity.** The ECS shows multiphasic kinetics that rely on activities of both photosystems, the cytochrome $b_6f$ complex, and ATP synthase. Averaged measurements of (**a**) ATeam and (**b**) Clover transformants are shown (biological replicates $N = 3 \pm$ SD).

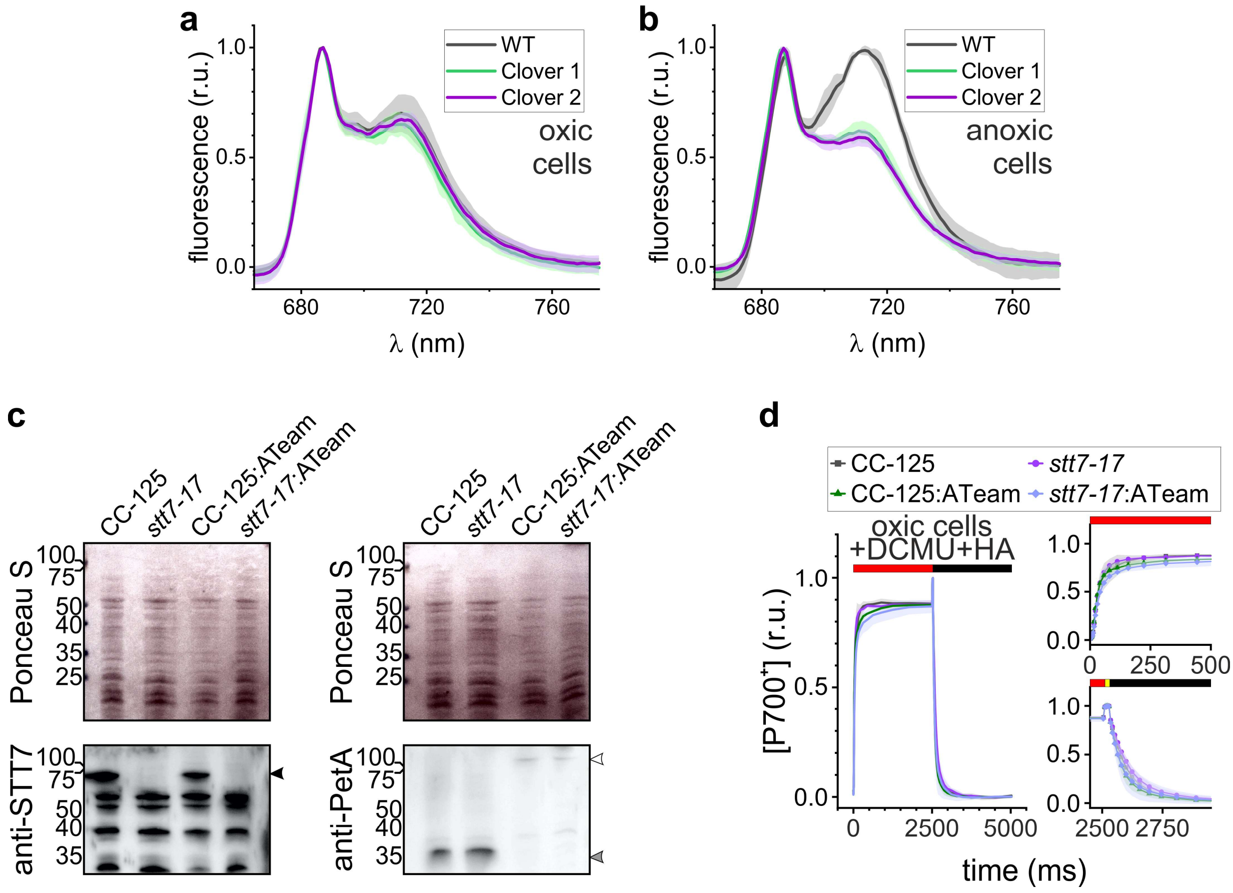


**Extended Data Fig. 6. State transition measurements in PetA-Clover fusion strains and characterization of PetA-ATeam fusion strains in the absence of STT7 kinase.** (**a**, **b**) 77 K chlorophyll fluorescence emission at ~712 nm, highlight the differences in antenna attached to photosystem (PS) I under State 1 (oxic) and State 2 (anoxic). These spectra were averaged and normalized to PSII-attached antenna emission signals at ~687 nm (biological replicates $N = 3 \pm$ SD). (**c**) Immunodetection of STT7 gene lesions (black arrowhead) and the ATeam-induced migration shift of PetA (grey arrowhead), yielding a fusion protein at ~100 kDa (white arrowhead). (**d**) Averaged P700 redox kinetics in PetA-ATeam fusion strains obtained in *stt7-17* and its reference CC-125 ($N = 3$ independent biological replicates, each measured in duplicate; mean ± SD). Actinic light, saturating pulse, and darkness are indicated via red, yellow, and black bars, respectively. For the double mutant kinetics, the amplitudes of PSI donor- and acceptor-side limitation, as well as the quantum yield, are labelled. PSII was inhibited by hydroxyl amine (HA) and 3-(3,4-Dichlorophenyl)-1,1-dimethylurea (DCMU).

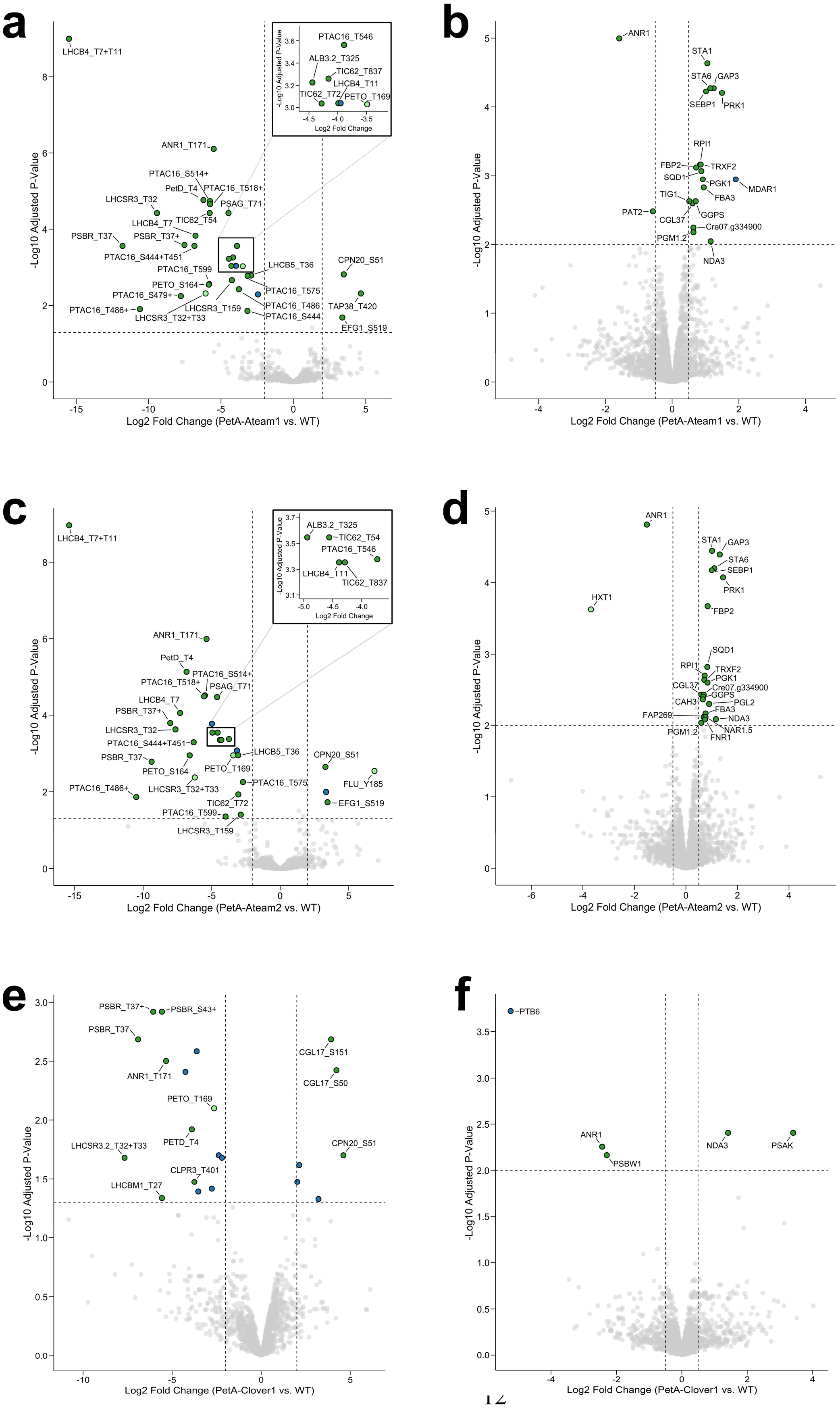
a
b
c
d
e
f
-Log10 Adjusted P-Value
Log2 Fold Change (PetA-Ateam1 vs. WT)
Log2 Fold Change (PetA-Ateam2 vs. WT)
Log2 Fold Change (PetA-Clover1 vs. WT)
Log2 Fold Change
LHCB4_T7+T11
ANR1_T171
PTAC16_S514+
PetD_T4
PTAC16_T518+
LHCSR3_T32
PSAG_T71
LHCB4_T7
TIC62_T54
PSBR_T37
PSBR_T37+
PTAC16_S444+T451
PTAC16_T599
PETO_S164
PTAC16_S479+
PTAC16_T486+
LHCSR3_T32+T33
LHCSR3_T159
LHCB5_T36
PTAC16_T575
PTAC16_T486
PTAC16_S444
CPN20_S51
TAP38_T420
EFG1_S519
PTAC16_T546
ALB3.2_T325
TIC62_T837
LHCB4_T11
TIC62_T72
PETO_T169
ANR1
STA1
STA6
GAP3
SEBP1
PRK1
RPI1
FBP2
TRXF2
SQD1
PGK1
MDAR1
FBA3
TIG1
PAT2
CGL37
GGPS
Cre07.g334900
PGM1.2
NDA3
HXT1
CAH3
PGL2
FAP269
NAR1.5
FNR1
FLU_Y185
PSBR_S43+
CGL17_S151
CGL17_S50
PETD_T4
LHCSR3.2_T32+T33
CLPR3_T401
LHCBM1_T27
PTB6
PSBW1
PSAK

**Extended Data Fig. 7. Volcano plots visualizing differences in protein phosphorylation (a, c, d) and protein abundances (b, d, f).** The respective panels show WT comparisons with PetA-Ateam 1 (**a, b**), PetA-Ateam 2 (**c, d**), and PetA-Clover 1 (**e, f**). Significant phosphorylation sites (FDR-adjusted p-value < 0.05, $|\log_2$ fold change$| > 2$) are highlighted in green (chloroplast-localized proteins) or blue (non-chloroplast proteins), with dashed lines marking thresholds. For proteins, significance thresholds are 0.01 (FDR-adjusted p-value) and 0.5 ($|\log_2$ fold change$|$). Data points shown in bright green represent instances where all values for a particular strain in the fold-change calculation had to be imputed due to undetectable peptide levels. Two positions linked via "+" (e.g., T32+T33) indicates concurrent phosphorylation at two sites, while a position followed by "+" alone denotes that the second site could not be unambiguously localized or multiple secondary sites exist on the same peptide. Phosphorylation fold-changes are normalized to protein abundance to account for expression differences.

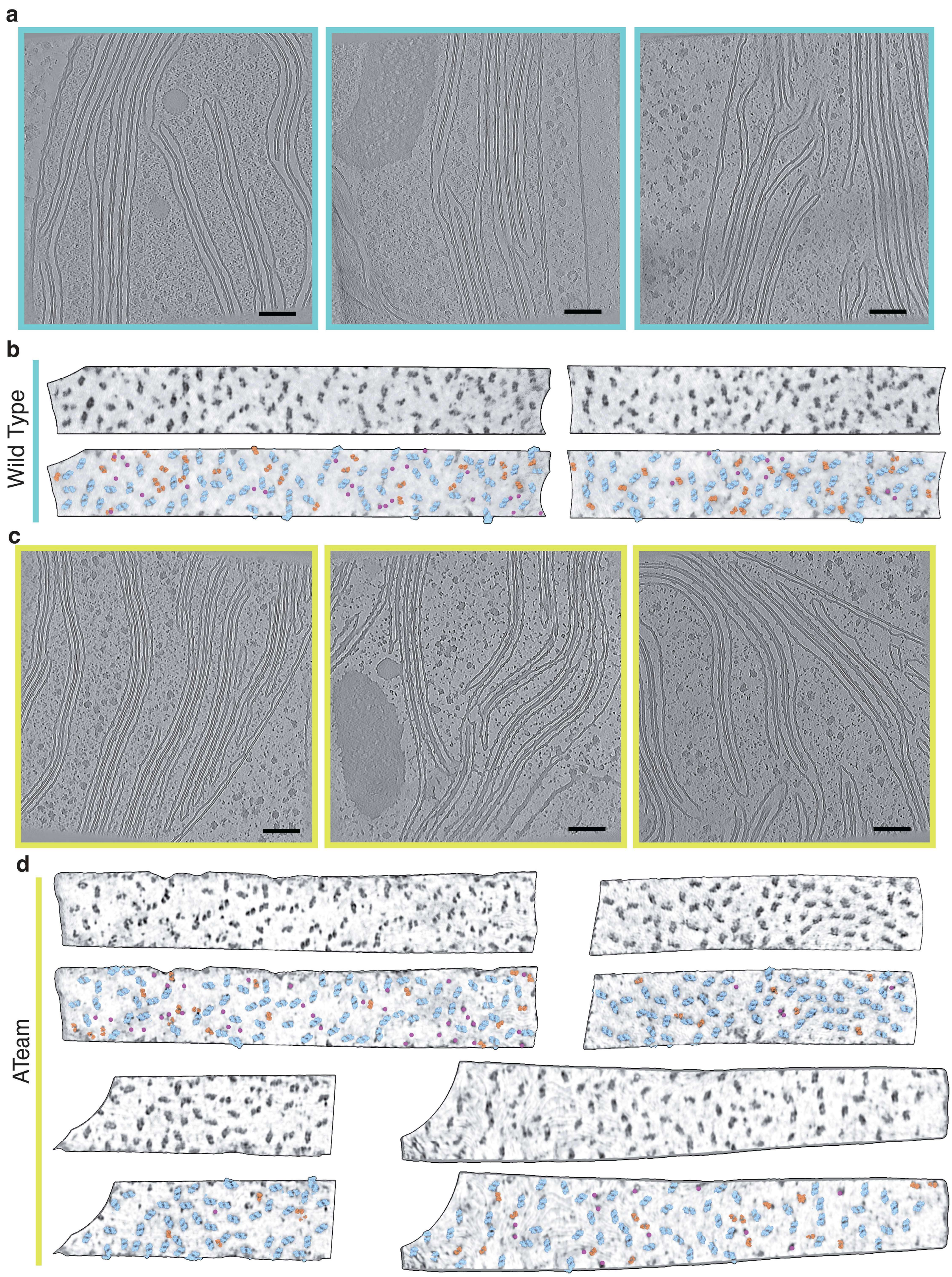


**Extended Data Fig. 8. Additional cryo-electron tomograms and representative membrane segments used for quantitative analysis.** (**a, b**) Representative cryo-electron tomographic *XY*

slices from three independent wild-type (**a**) and PetA-ATeam (**c**) cells showing appressed thylakoid membrane regions used for particle analysis. (**b, d**) Representative appressed thylakoid membrane segments extracted from WT and PetA-ATeam tomograms, respectively, and used for quantitative particle identification. Top, intensity projections of the raw tomograms across the membrane thickness. Bottom, corresponding membrane segments with manually assigned particles overlaid: photosystem II (PSII, blue), cytochrome $b_6f$ (orange), and unassigned particles (magenta).

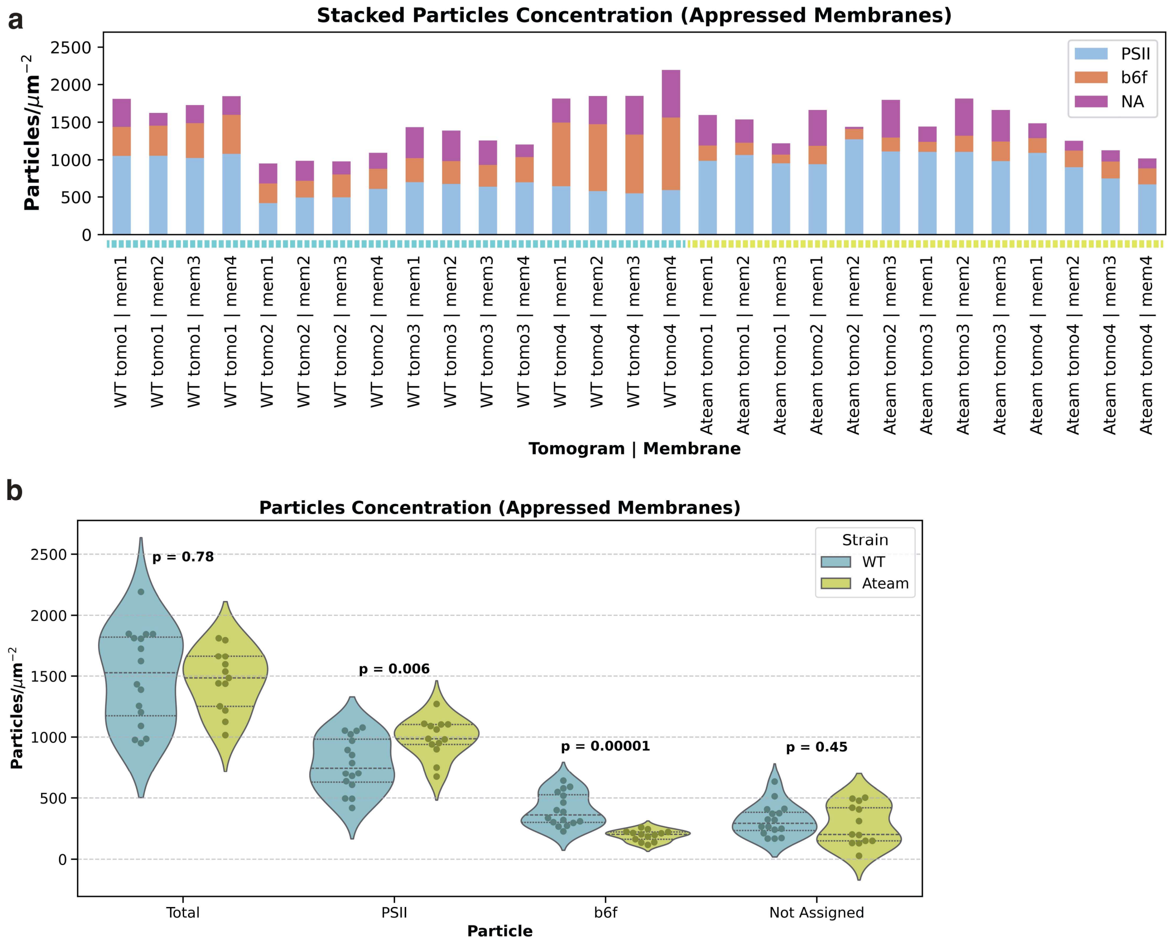


**Extended Data Fig. 9. Quantitative analysis of particle concentrations in appressed thylakoid membranes.** (**a**) Stacked bar plots showing particle concentrations measured in individual appressed thylakoid membrane segments from wild-type (WT) and PetA-ATeam (Ateam) tomograms. Bars represent total particle concentration per membrane segment, decomposed into contributions from photosystem II (PSII, blue), cytochrome $b_6f$ (orange), and unassigned particles (NA, magenta). Each bar corresponds to one analyzed membrane segment, grouped by tomogram and strain as indicated along the x-axis. (**b**) Violin plots summarizing particle concentrations in appressed thylakoid membranes from WT (cyan) and PetA-ATeam (yellow) cells. Distributions are shown for total particle concentation, PSII, cytochrome $b_6f$, and unassigned particles. Individual data points represent single membrane segments. Horizontal dashed lines indicate median and interquartile ranges. P values are shown for comparisons between strains (two-sided statistical test), highlighting a significant reduction in cytochrome $b_6f$ density and a modest increase in PSII density in PetA-ATeam membranes, while total particle concentration and unassigned particles remain unchanged.

**Extended Data Tab. 1. Primers, synthesized genes and mass spectrometric results.**

Download as separate spreadsheet file
