## Supplementary figures and images for "Lateral organization of cytochrome *b_6_f* in thylakoid membranes controls photosynthetic electron transfer efficiency"

### Supplemental Figure 1

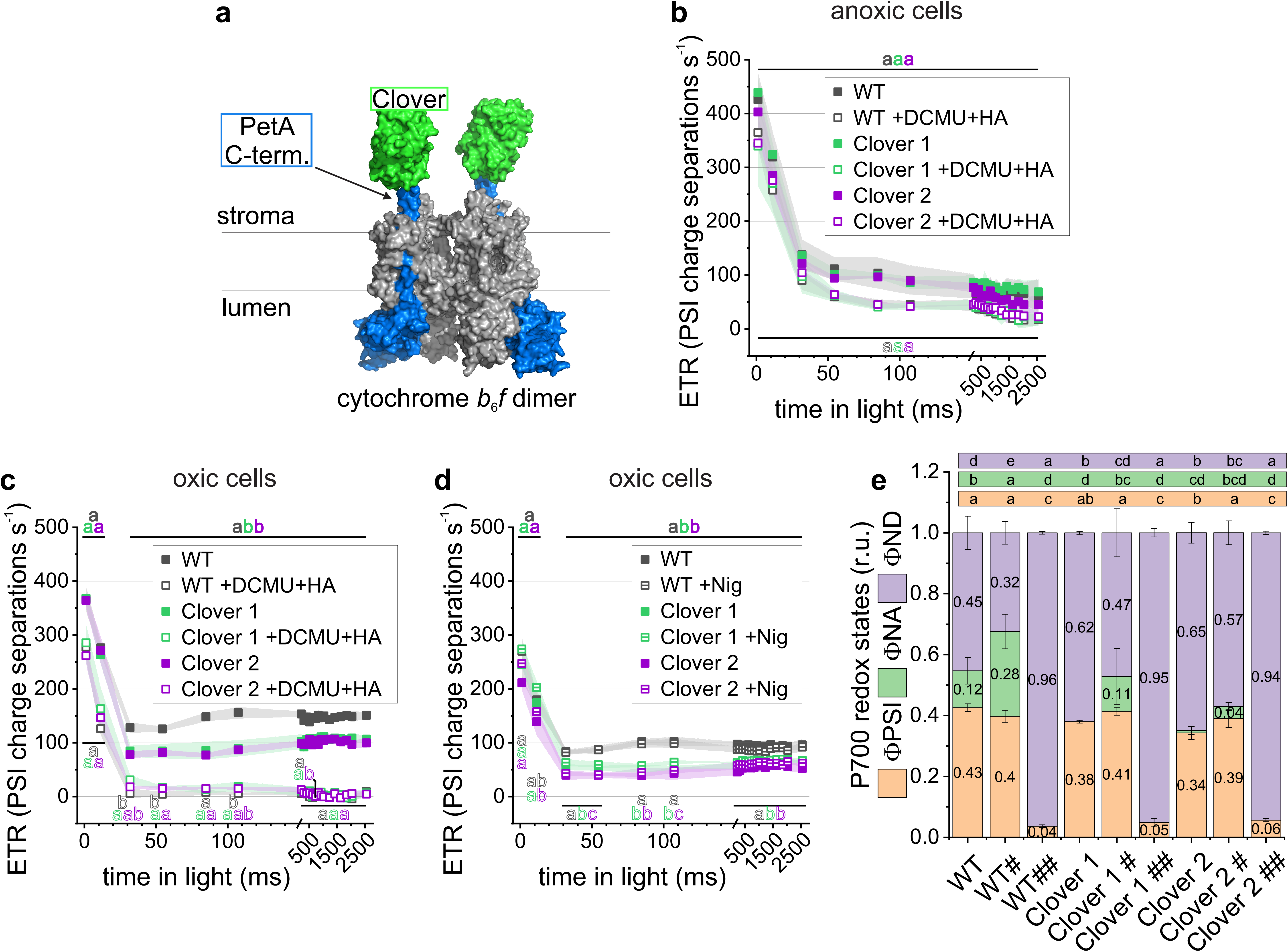

### Supplemental Figure 2

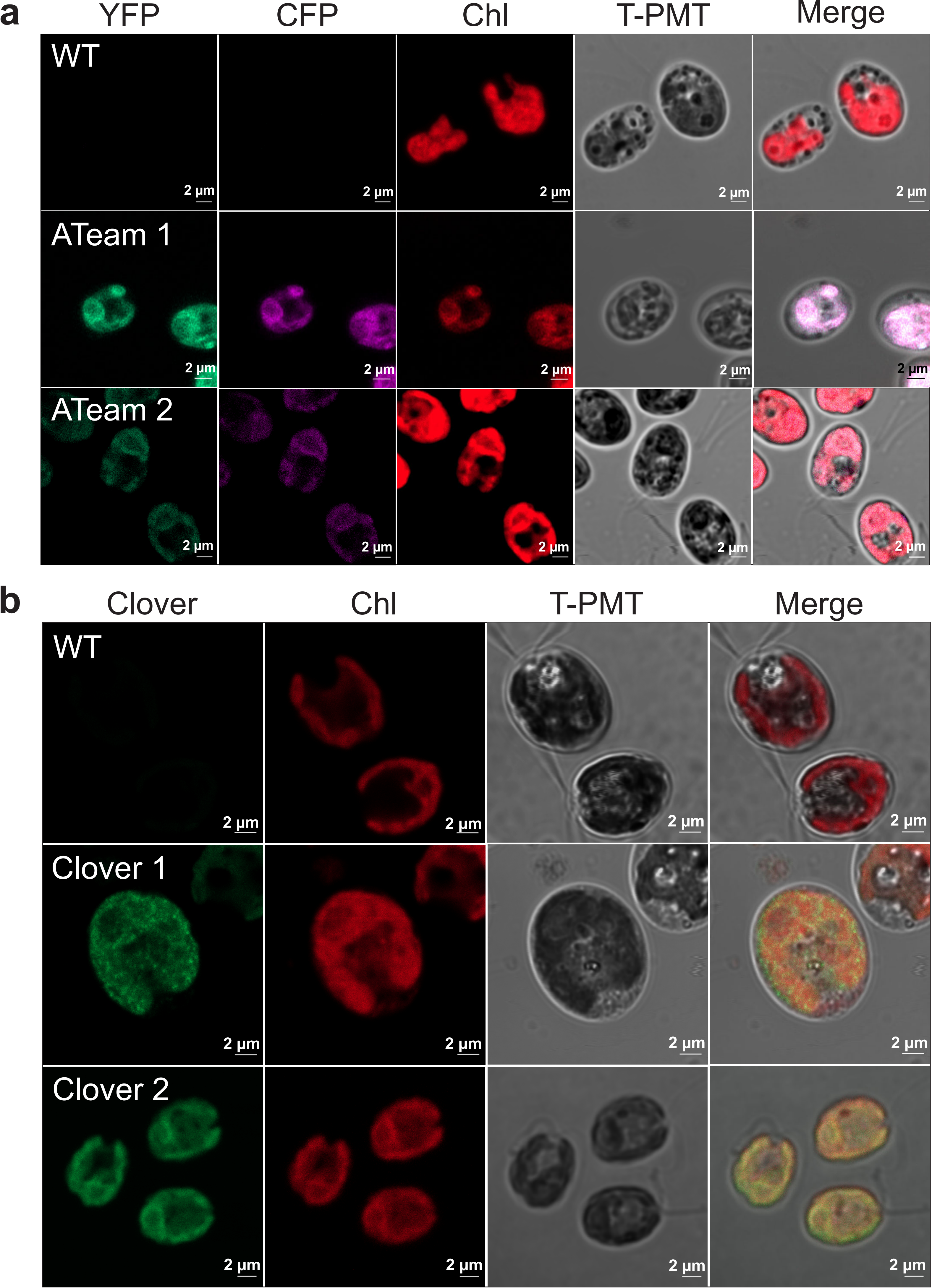

### Supplemental Figure 3

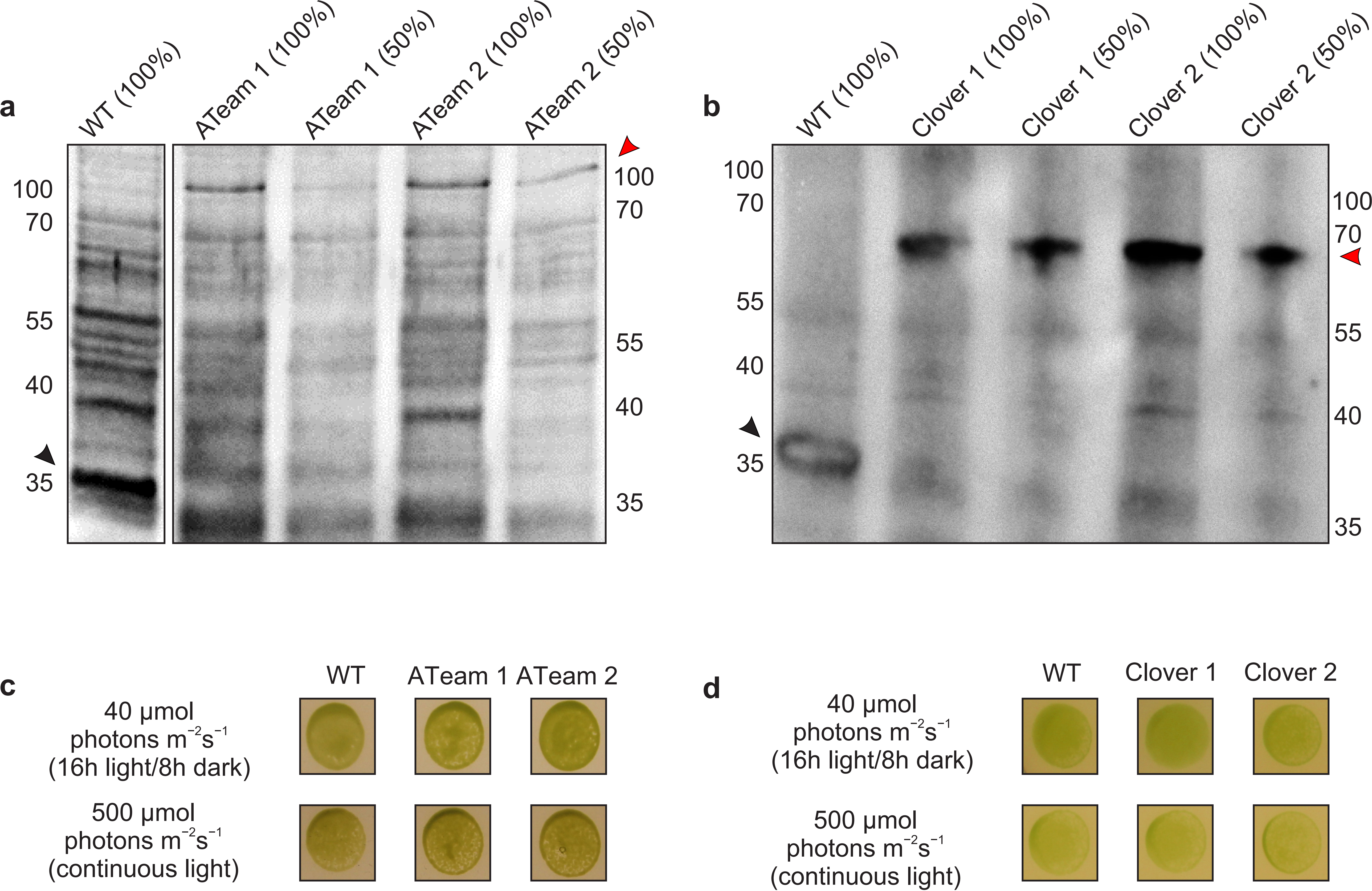

### Supplemental Figure 4

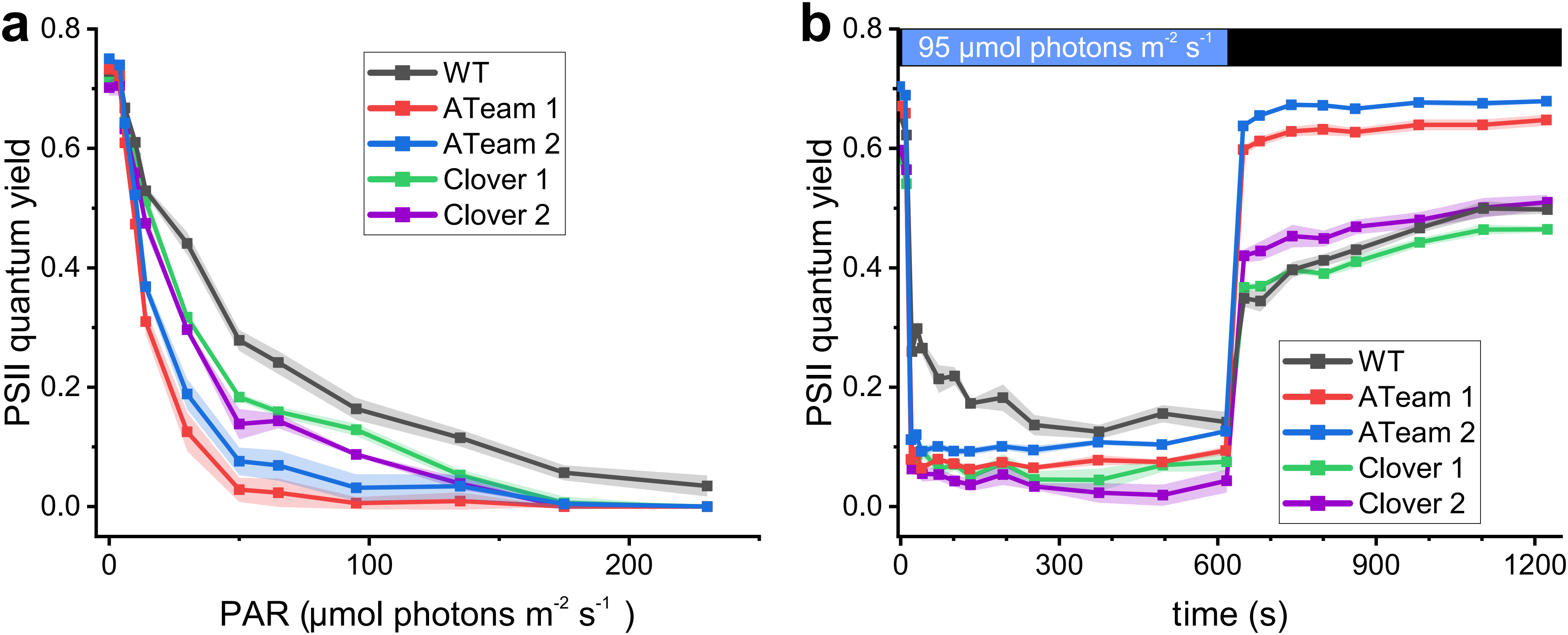

### Supplemental Figure 5

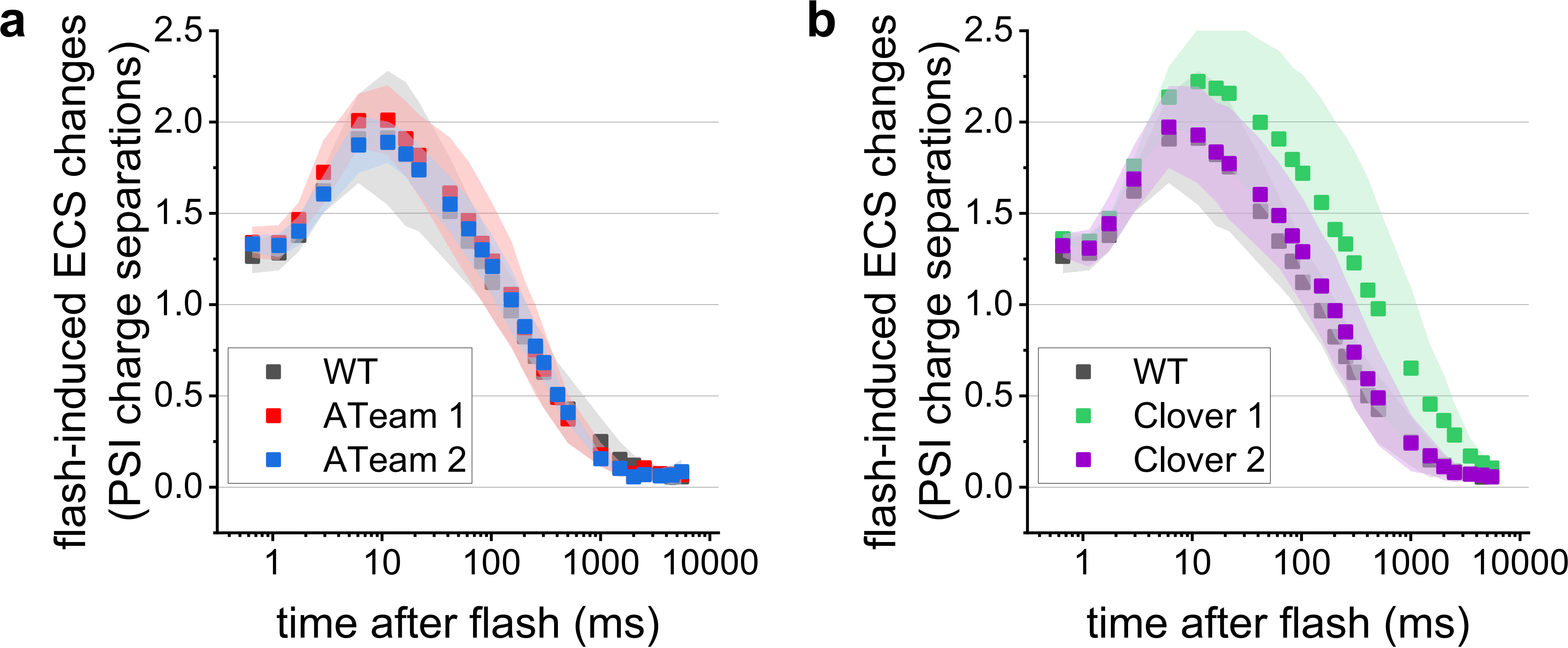

### Supplemental Figure 6

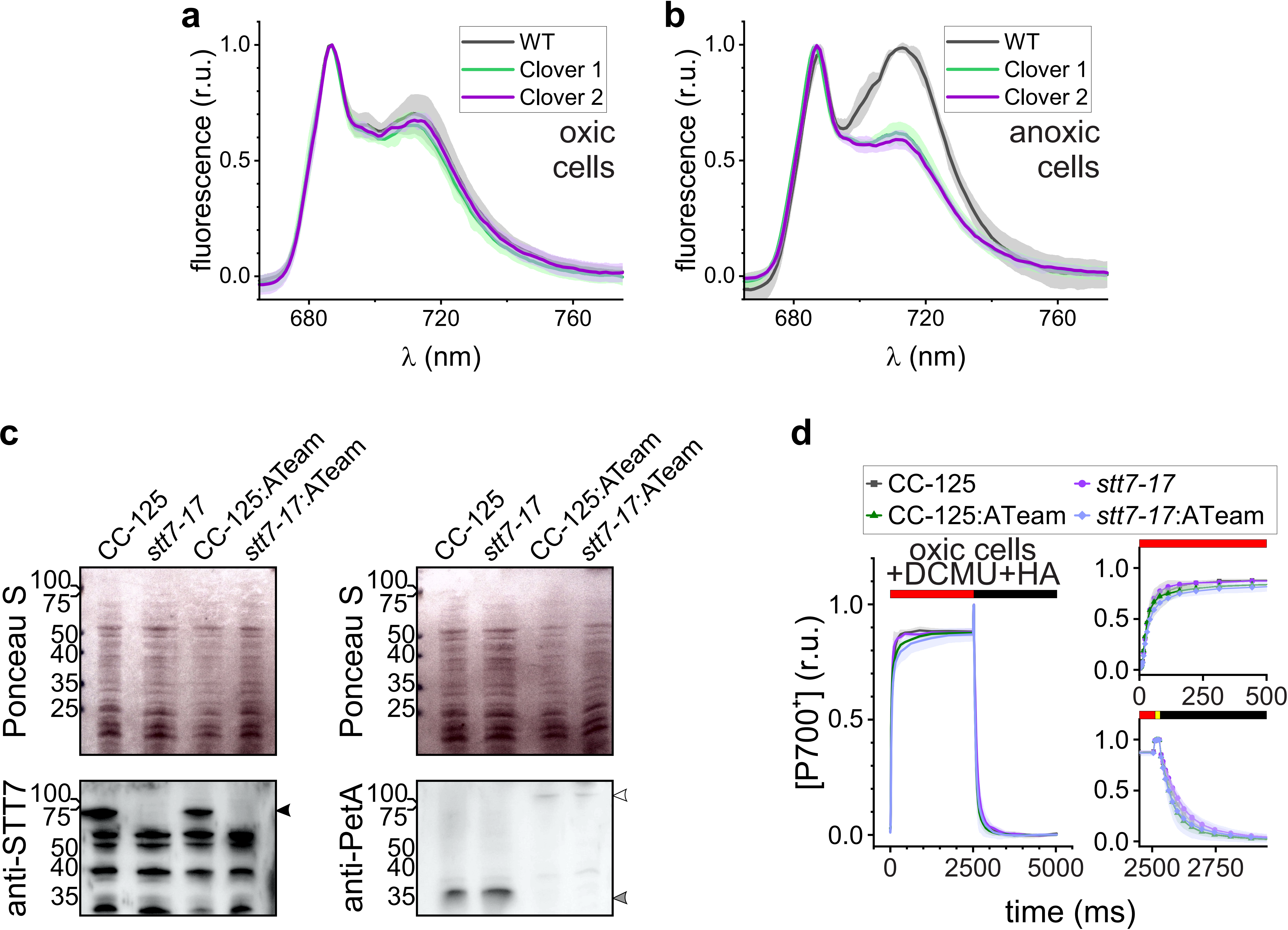

### Supplemental Figure 7

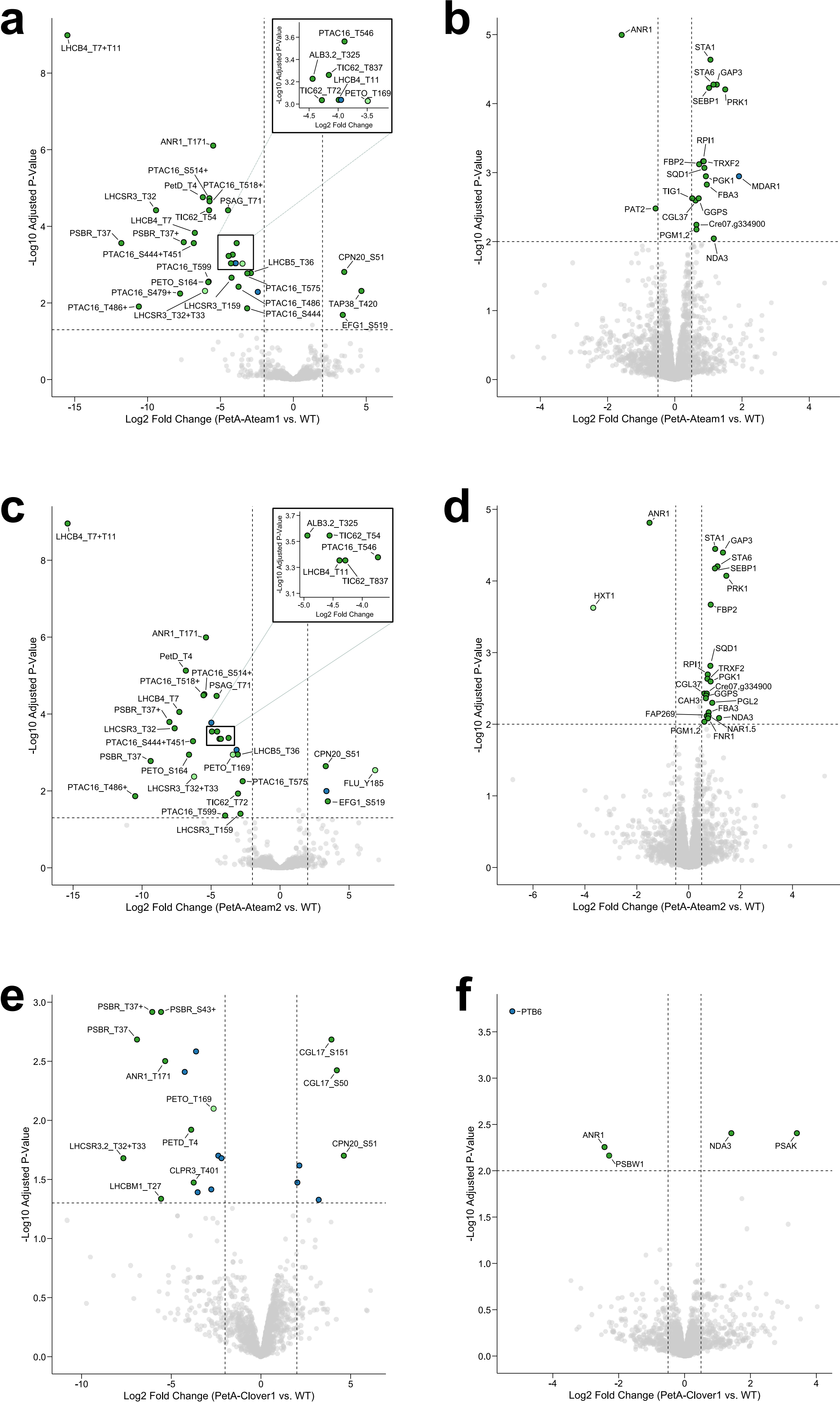

### Supplemental Figure 8

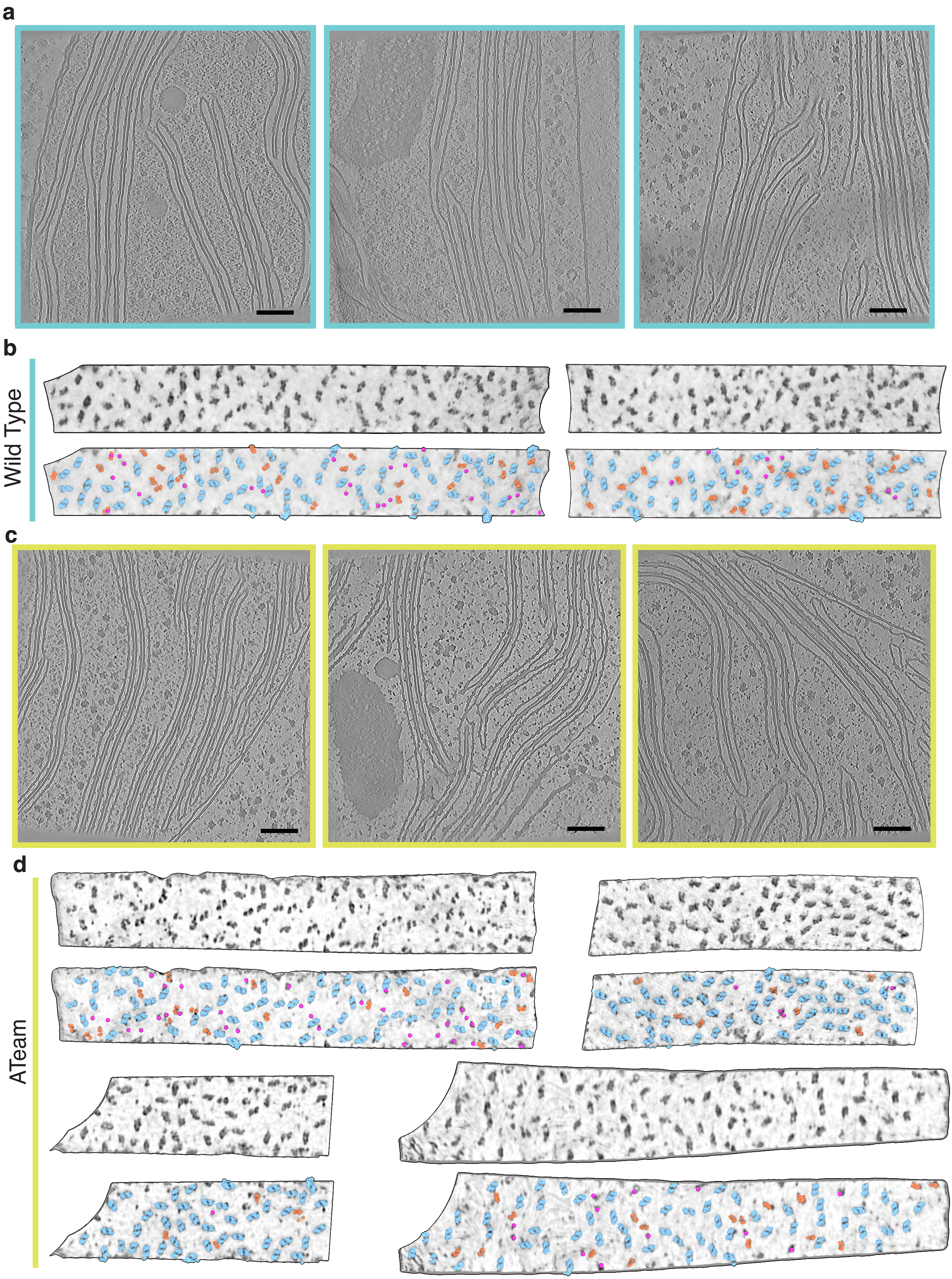

### Supplemental Figure 9

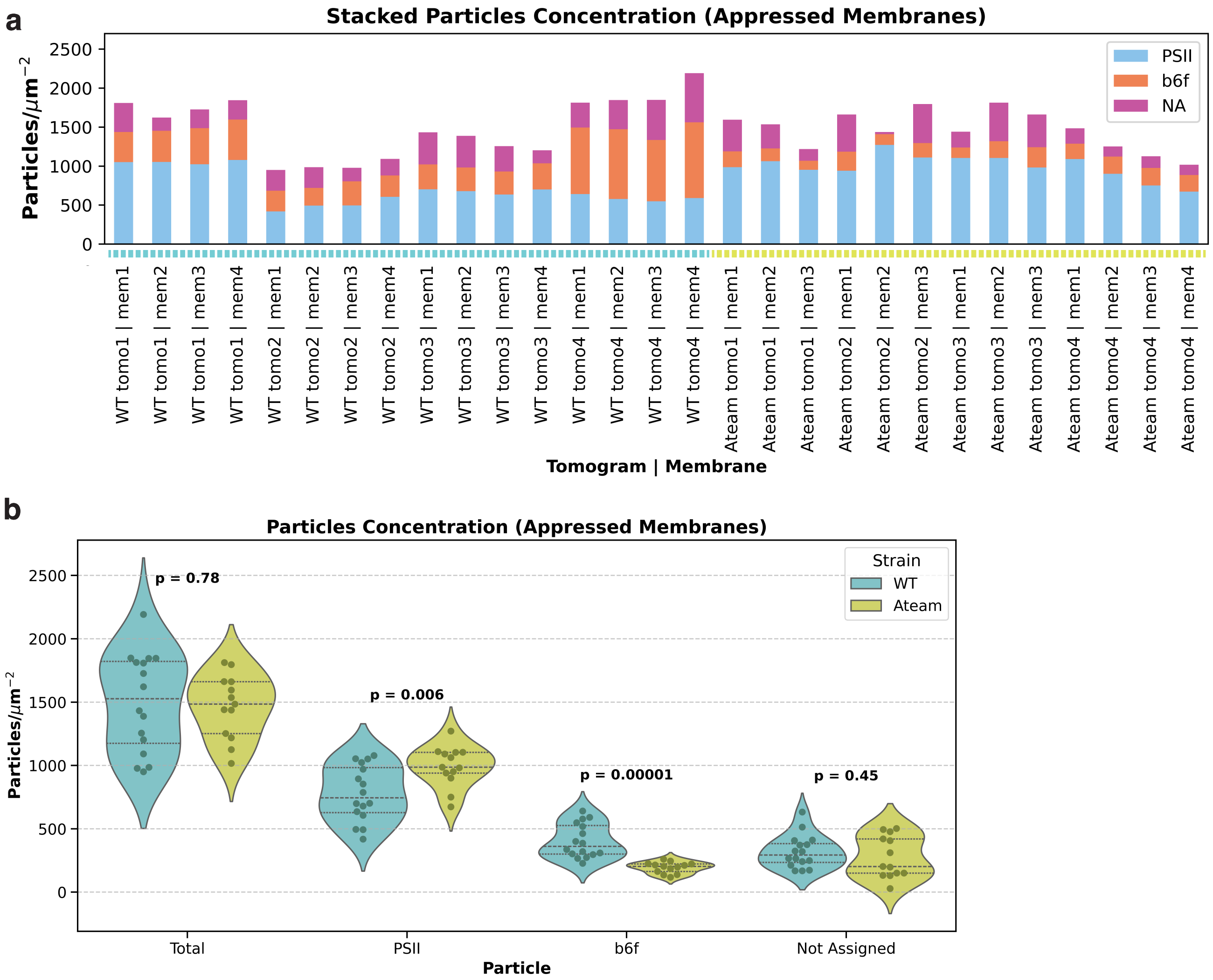
